# SPTF-3/SP1 orchestrates mitochondrial biogenesis upon ribosomal stress and acute starvation

**DOI:** 10.1101/2023.08.18.553493

**Authors:** Johannes CW Hemerling, Victor Pavlenko, Marija Herholz, Linda Baumann, Aleksandra Zečić, Alexandra Kukat, Milica Popovic, Karolina Szczepanowska, Estela Cepeda Cores, David Vilchez, Leo Kurian, Aleksandra Trifunovic

## Abstract

When cells have increased energy demand, they respond by elevating the production of new mitochondria through the process of mitochondrial biogenesis. This complex physiological undertaking requires precise coordination of mitochondrial and nuclear gene expression to extend the existing mitochondrial network in the cell. Using C. elegans as a model system we have identified stress-induced transcription factor SPTF-3 as a novel regulator of mitochondrial biogenesis on-demand upon increased heat stress, dietary restriction, and acute starvation. We show that SPTF-3 also regulates ATFS-1, the main transcriptional regulator of UPR^mt^ (mitochondrial unfolded protein response). Thus, by orchestrating two parallel programs – mitochondrial biogenesis and UPR^mt^, SPTF-3 safeguards mitochondrial wellbeing and function upon stress, thus allowing survival in unfavorable conditions. Mitochondrial biogenesis is induced by disturbances in cytoplasmic ribosomal assembly, which leads to preferential translation of SPTF-3. Importantly, we demonstrated that the role of SPTF-3 in the regulation of mitochondrial biogenesis upon nutrient deprivation is conserved in mammals through its homolog SP1.

## Introduction

Mitochondrial mass, organization, and function vary between different cell types and are dynamically adapted to environmental and intracellular stimuli. An increase in mitochondrial biogenesis is often the first response to high energy demands in physiological and pathological conditions. This is a highly coordinated growth of pre-existing mitochondria that requires transcriptional activation, synthesis and import of mitochondrial proteins; delivery of lipids to mitochondrial membranes, and corresponding activation of mtDNA replication and organelle gene expression. As a relic of the bacterial origin, mitochondria possess their own DNA, mitochondrial DNA (mtDNA) that encodes only a handful of subunits of the electron transport chain (ETC). The rest of the over 1000 proteins that form mitochondria, including all the proteins that mediate mtDNA replication, transcription, and translation are encoded in the nuclear genome, synthesized in the cytoplasm, and imported into the mitochondria. Thus, mitochondrial biogenesis and remodeling are extremely complex and rely on coordinated regulation of both nuclear and mtDNA gene expression.

Considerable progress has been made in deciphering mitochondrial biogenesis-related proteins and genes that function in health and pathology-related circumstances. Several transcription factors (TFs) and co-factors have been described to control nuclear (n) DNA-encoded mitochondrial proteins^1, 2^. Nuclear respiratory factor 1 (NRF1) and 2 (NRF2)-binding sites can frequently be found in the proximal promoters of many mammalian mitochondrial genes, including many OXPHOS subunits and genes involved in mitochondrial transcription, offering a mechanism for bi-genomic transcription control^2^. Peroxisome-proliferator activated receptor α (PPARα) was the first nuclear receptor shown to regulate mitochondrial metabolism through transcriptional control of mitochondrial FAO enzymes^3^. The second family of nuclear receptors involved in the regulation of mitochondrial biogenesis consists of estrogen-related receptors (ERR). Although initially suggested to activate the expression of FAO enzymes in oxidative tissues, later studies revealed that ERRs regulate the expression of genes involved in effectively all mitochondrial energy-producing pathways^4^.

The discovery of the peroxisome proliferator-activated receptor gamma co-activator 1 (PGC-1) family of co-activators, consisting of PGC-1α, PGC-1ß, and PGC-related coactivator (PRC) has offered a possible mechanism for the integration of physiological stimuli and orchestration of transcription activity in mammalian mitochondrial biogenesis^5^. PGC-1 family members bind to the respective TFs, thereby being recruited to the transcription site of the target gene. The critical feature of these coactivators is their high versatility in interacting with distinct TFs, including NRF-1/2, PPARs, and ERRs. This allows them to activate different biological programs in a tissue-specific manner^5^. In mitochondria-rich tissues like the heart, brown fat, and muscles, expression of PGC-1α is strongly induced upon cold exposure and short-term exercise, resulting in increased expression of mitochondrial genes^6–8^. However, the role of these co-activators in the regulation of mitochondrial biogenesis outside mammals is ambiguous, as it seems that they (PGC-1α) do not play a role in the regulation of mitochondrial content in lower vertebrates^9^, or are completely absent in invertebrates.

The pathways that sustain mitochondrial biogenesis through aging, such as caloric restriction or exercise, seem to increase fitness and longevity in multiple species^10, 11^. Remarkably, transcriptional regulators of mitochondrial biogenesis described so far are specific for vertebrates and no homologs could be found in invertebrates even though mitochondrial biogenesis on-demand is well-documented in different stress conditions across various fila^11^. Therefore, it is crucial to identify novel mechanisms and factors that regulate mitochondrial biogenesis on-demand to understand the fine-tuned regulation and the downstream metabolic effects.

We performed a genome-wide screen for factors that promote mitochondrial biogenesis in *Caenorhabditis elegans* (*C. elegans*) and identified perturbation in the levels and assembly of cytoplasmic ribosomes as a major pathway promoting mitochondrial gene expression. Depletion of individual ribosomal proteins leads to inefficient ribosomal biogenesis resulting in altered translation fidelity. This induces the synthesis of SPTF-3, a homolog of the mammalian SP1 transcription factor, that in turn promotes mitochondrial gene expression and remodels the mitochondrial network. We further show that in worms this pathway is activated upon starvation, caloric restriction, and heat stress and is highly conserved in mammalian cells, where it is initiated upon nutrient deprivation.

## Results

### Cytosolic ribosomal proteins suppress mitochondrial biogenesis

To identify novel factors regulating mitochondrial biogenesis on-demand we used *C. elegans* as a model system. To this end, we generated a transgenic strain expressing GFP under the control of the *mtss-1* promoter. MTSSB-1 is a homolog of mammalian mitochondrial single-strand binding protein (SSBP1), whose expression directly correlates with mtDNA levels, reflecting the changes in mitochondrial mass^12^. When grown at a standard maintenance temperature of 20°C, GFP in the reporter stain (*pmtss-1::gfp*) was expressed predominantly in the pharynx and the posterior part of the worm. We and others previously reported that shifting worms to a higher temperature (25°C) significantly increases the amount of mtDNA and mitochondria in somatic tissues^13, 14^. Indeed, after shifting worms to 25°C, *pmtss-1::gfp* expression was robustly upregulated in the intestine (Fig. 1A). The upregulation of the fluorescent signal was confirmed by Flow Cytometry Analyses (FACS) (Extended Data Fig. 1A). In agreement, animals grown on 25°C showed increased expression of endogenous *mtss-1* (Fig. 1B). This was accompanied by an increase in the mtDNA copy number and oxygen consumption (Extended Data Fig. 1B-C). Changes in *mtss-1* expression are independent of the mitochondrial biogenesis in the germ line, as we have previously shown that mtDNA increases on 25°C stems from the somatic tissues as is observed also in gonad-less *glp-4(bn2)* mutant^13^. Thus, the *pmtss-1::gfp* reporter that we developed could be used in a further screen for TFs regulating somatic mitochondrial biogenesis on-demand.

**Figure 1.**
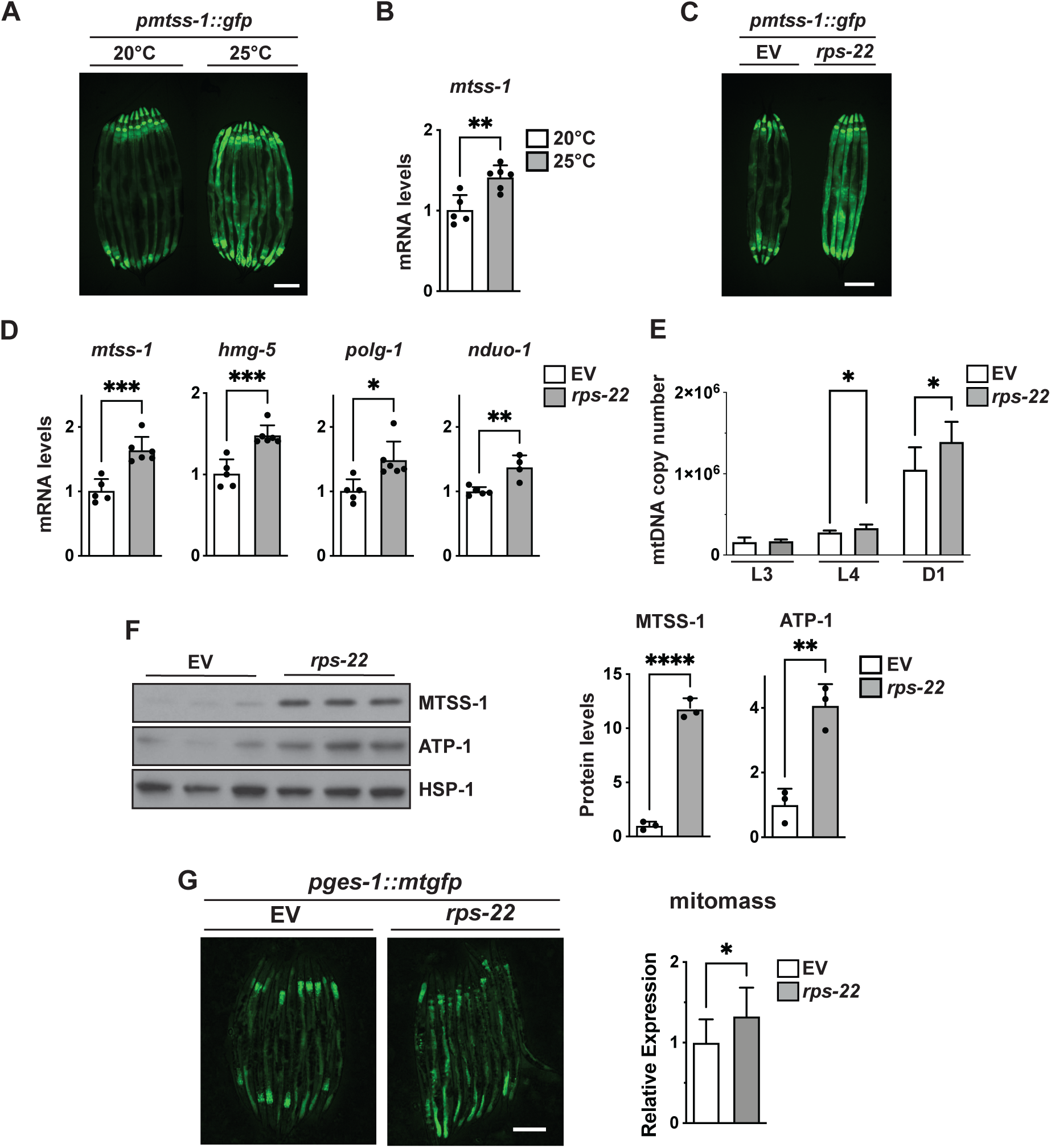
Depletion of cytosolic ribosomal subunits induces mitochondrial biogenesis. **A)** Fluorescent images of a strain expressing *gfp* under *mtss-1* promoter (*pmtss-1::gfp*) grown on 20°C and 25°C; **B)** The expression levels of *mtss-1/SSBP-1* in worms grown on 20°C and 25°C; **C)** Fluorescent images of *pmtss-1::gfp* reporter strain upon *rps-22* RNAi; **D**) Steady-state transcript levels of *mtss-1/SSPB1, hmg-5/TFAM, polg-1/POLG* and *nduo-1/mt-ND1* levels upon *rps-22* RNAi; **E)** mtDNA copy number measured by qPCR in L3, L4 and adult day 1 (D1) animals upon *rps-22* RNAi; **F)** Steady-state levels of mitochondrial proteins MTSS-1/SSBP1 and ATP-1/ATP5A upon *rps-22* RNAi; **G)** Total mitomass measured using a reporter with mitochondria-targeted *gfp* expression in the gut (*pges-1::mtgfp*). Representative florescent images (*left*) and quantitative analysis (*right*). Data are presented as mean ± SD. \**p*<0.05, \*\**p*<0.01, \*\*\**p*<0.001, \*\*\*\**p*<0.0001, unpaired t-test.

To understand more about the regulation of mitochondrial biogenesis, we started a genome-wide RNAi screen for suppressors of mitochondrial biogenesis in *C. elegans* from the Ahringer library^15^. After analyzing the results obtained using knockdown of genes on chromosome III, we observed strong GFP induction when genes encoding multiple cytosolic ribosomal proteins were silenced (Fig. 1C, Extended Data Fig. 1D and Table 1), hence decided to investigate this in more detail, using *rps-22* silencing as a model for further analyses. The increase in *mtss-1* expression was mirrored by higher levels of *hmg-5*/*TFAM*, *polg-1*/*POLG* (mtDNA polymerase), and *nduo-1*/*MT-ND1* (mtDNA-encoded NADH-ubiquinone oxidoreductase chain 1) transcripts (Fig. 1D). Mitochondrial biogenesis in the somatic tissues of *C. elegans* predominantly occurs between the larval stages L3 and late L4^13^. Already at the L4 stage, the mtDNA copy number was increased upon *rps-22* knockdown (Fig. 1E) coinciding with the high expression from the *mtss-1* promoter (Extended Data Fig. 1E). This increased steady-state levels of ATP-1/ATP5A, an ATP synthase subunit, and MTSS-1 (Fig. 1F and Extended Data Fig. 1F). Using a reporter with mitochondria-targeted GFP expression in the gut (*pges-1::mtgfp*) we could also demonstrate overall higher mitochondrial mass upon *rps-22* depletion (Fig. 1G). Collectively, these data suggest that ribosomal dysfunction leads to an increase in mitochondrial biogenesis.

**Table 1.** RNAi clones from Ahringer or Vidal RNAi library.

| **Name** | **Sequence** | **Ahringer RNAi library** |
| --- | --- | --- |
| *atfs-1* | ZC376.7 | Vidal RNAi library |
| *cep-1* | F52B5.5 | I-4G17 |
| *daf-15* | C10C5.6 | IV-4H20 |
| *gei-17* | W10D5.3 | I-4D09 |
| *ife-2* | R04A9.4 | X-1A21 |
| *let-363* | B0261.2 | I-9A17 |
| *mlst-8* | C10H11.8 | I-2I05 |
| *ncbp-1* | F37E3.1 | I-3G09 |
| *ncbp-2* | F26A3.2 | I-3D16 |
| *rict-1* | F29C12.3 | II-8J07 |
| *rpl-26* | F28C6.7 | II-6C16 |
| *rpl-35* | ZK652.4 | III-4I21 |
| *rpoa-2* | F14B4.3 | I-4H13 |
| *rps-6* | Y71A12B.1 | I-9F06 |
| *rps-7* | ZC434.2 | I-5M09 |
| *rps-22* | F53A3.3 | III-1O10 |
| *rict-1* | F29C12.3 | II-8J07 |
| *rsks-1* | Y47D3A.16 | III-8C23 |
| *sinh-1* | Y57A10A.20 | Vidal RNAi library |
| *spg-7* | Y47G6A.10 | Vidal RNAi library |
| *sptf-3* | Y40B1a.4 | I-6F02 |
| *W04D2.4* | W04D2.4 | V-14I24 |

### Mitochondrial biogenesis is regulated by the loss of ribosomal subunits not a general decrease in translation

We next tested if depletion of other genes encoding ribosomal proteins or proteins involved in translation would likewise induce *mtss-1–*driven GFP expression. We performed a targeted screen using the *pmtss-1::gfp* reporter strain and all available RNAi clones covering the entire spectrum of cytosolic ribosomes, translation initiation, elongation and termination factors, and other factors involved in protein synthesis, e.g. tRNA synthetases (Extended Data Table 2). Strikingly, 85% of all silenced ribosomal proteins strongly induced *pmtss-1::gfp* expression regardless if the small or large ribosomal subunits were targeted (Fig.1C, Extended Data Fig. 1D-1E and Table 2). As ribosomal subunits are essential for development, the depletion of many caused developmental delay or arrest, resulting in a smaller worm size (Extended Data Fig. 2A), and, for the ones that reached adulthood, greatly reduced brood size. To overcome the developmental arrest, we exposed worms to diluted RNAi that still provided strong induction of GFP expression, but fewer side effects, and used this condition for all further experiments. In contrast, depletion of translation factors failed to alter GFP expression, suggesting that they could either compensate for each other’s loss or that the initial signal for increased mitochondrial biogenesis originates from a defect in ribosomal assembly, rather than protein synthesis. To further investigate this, we downregulated the expression of genes encoding the translation initiation factor IFE-2/eIF4E and GCN-2/GCN2, a kinase that regulates the activity of eukaryotic translation initiation factor 2 alpha/eIF2a and was previously shown to be a longevity assurance factor for mitochondrial mutants^16^. Interestingly, in both mutants, the signal from the *pmtss-1::gfp* seemed even lower than control (N2) (Fig. 2A). We also did not observe change in mtDNA copy number in the *ife-2(ok306)* mutant (Fig. 2B). Nevertheless, *rps-22* depletion in *gcn-2(ok871)* and *ife-2(ok306)* mutants increased *mtss-1* expression (Fig. 2C and Extended Data Fig. 2B), again suggesting diverse pathways. These data imply that defect in ribosomal biogenesis, as demonstrated by the changes in ribosomal profiles (Fig. 2D), rather than repression of translation, leads to increased mitochondrial biogenesis.

**Figure 2.**
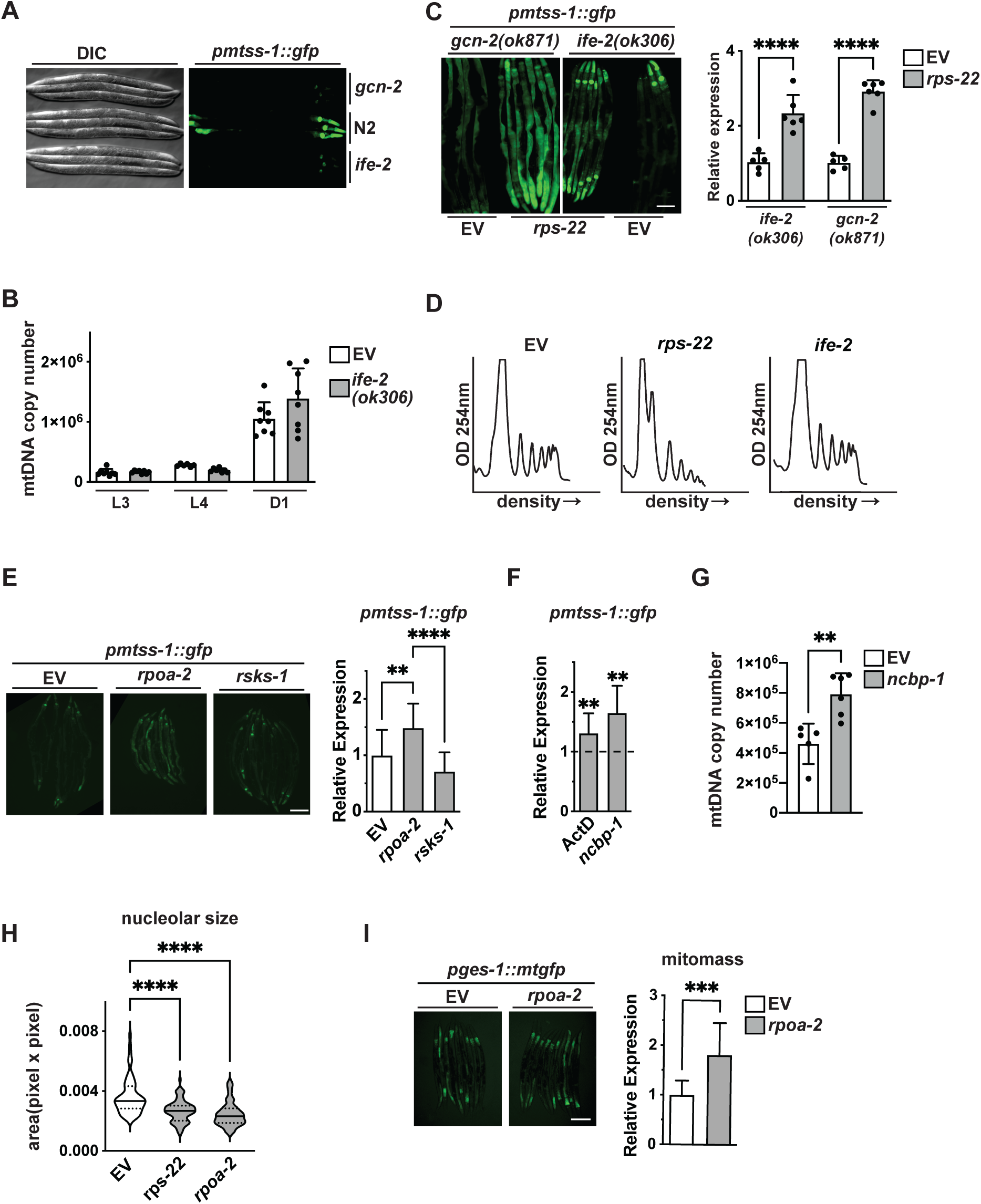
Ribosomal stress, not translation, induces increased MTSS-1. **A)** Fluorescent and control (DIC) images of *pmtss-1::gfp* reporter strain upon *gcn-2* and *ife-2* RNAi; **B)** mtDNA copy number measured by qPCR in L3, L4 and adult day 1 (D1) animals in control and *ife-2(ok306)* mutants; **C)** *mtss-1* expression levels measured using *pmtss-1::gfp* reporter in *gcn-2(ok871)* and *ife-2(ok306)* mutants upon *rps-22* RNAi. Representative florescent images (*left*) and quantitative analysis (*right*); **D)** Ribosomal profiling upon *rps-22* and *ife-2* RNAi; **E)** *mtss-1* expression levels measured using *pmtss-1::gfp* reporter upon *rpoa-2* and *rsks-1* RNAi. Representative florescent images (*left*) and quantitative analysis (*right*); **F)** *mtss-1* expression levels measured using *pmtss-1::gfp* reporter upon treatment with Actinomycin D (ActD) or *ncbp-1* RNAi; **G)** mtDNA copy number measured by qPCR in L3, L4 and adult day 1 (D1) animals upon *ncbp-1* RNAi; **H)** Nucleolar size analysed using quantification fluorescence of *pfig-1::fib-1::gfp* reporter upon *rpoa-2* RNAi. Representative images in Figure S2F; **I)** Total mitomass measured using a *pges-1::mtgfp* reporter upon *rpoa-2* RNAi. Quantitative data are presented as mean ± SD. \*\**p*<0.01, \*\*\**p*<0.001, \*\*\*\**p*<0.0001. Unpaired t-test was used in C, F, G, H. One-way ANOVA with Tukey post hoc test was used in B, E, I.

**Table 2.**
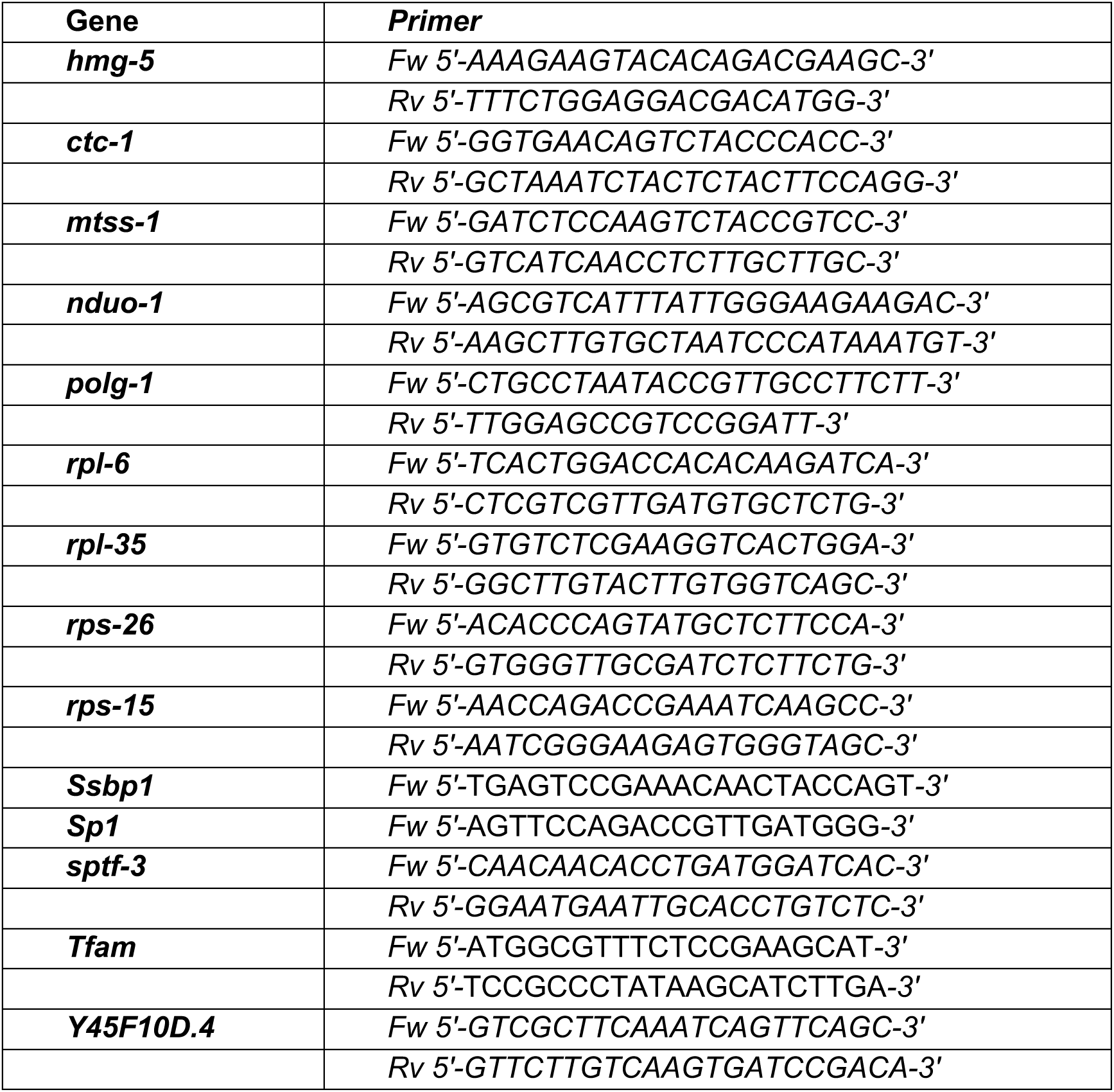
Primers.

Intrigued by these findings, we took a closer look at *C. elegans* TOR (*Ce*TOR) signaling as a potent modulator of cytosolic translation. We silenced different components of *Ce*TOR complexes (*let-363/MTOR* and *mlst-8/LST8*), including subunits specific for mTORC1 (*daf-15/RPTOR*) or mTORC2 (*rict-1/RICTOR* and *sinh-1/MAPKAP1*) complexes in nematodes (Extended Data Fig. 2C). This did not result in major changes of expression from *mtss-1* promoter, which could be strongly induced by simultaneous *rps-22* depletion (Extended Data Fig. 2C). Similarly, switch to higher temperature still induced the *mtss-1* expression in the mutant of *rsks-1* (Extended Data Fig. 2D), a homolog of S6 kinase, an extensively studied effector of the TORC1 complex^17^. Adding further evidence that mTOR does not play a direct role in the regulation of mitochondrial mass on-demand, the presence of *Ce*TOR components did not affect mitochondrial levels (Extended Data Fig. 2E).

In contrast to *rsks-1* loss, depletion of *rpoa-2/POLR1B*, a subunit of RNA polymerase I (RNAP I) that resides in the nucleolus and transcribes only ribosomal RNAs, thus accounting for over 50% of the total RNA synthesized in a cell, induces *mtss-1* expression (Fig. 2E). Preventing assembly of the 60S ribosome by depletion of the nuclear cap-binding protein 1 (*ncbp-1/NCBP1)* or treatment with Actinomycin D (ActD) also resulted in increased *mtss-1::gfp* expression, and higher mtDNA copy number (Fig. 2F-2G). Ribosomal biogenesis occurs in the nucleolus and is dependent on its architecture, thus nucleolar size relates to ribosomal stress^18^. We could show that both *rpoa-2/POLR1B* and *rps-22* depletion significantly decreases nucleolar size as measured by the expression of the GFP-tagged nucleolar protein FIB-1 (Fig. 2H and Extended Data Fig. 2F). These results clearly link nucleolar size and the defect in ribosomal production to enhanced mitochondrial biogenesis and consequently increased mito-mass (Fig. 2H-2I). Thus, ribosomal biogenesis stress seems to be the primary signal for increased mitochondrial biogenesis upon depletion of individual cytosolic ribosomal subunits.

Different types of cellular stresses were reported to activate P53 by blocking specific steps of ribosome biogenesis and inducing nucleolar stress^18, 19^. To investigate if P53 is involved in the regulation of mitochondrial biogenesis on-demand, we depleted *cep-1*, a *C. elegans* homolog of P53 under conditions that induce *mtss-1* expression (Extended Data Fig. 2G-2H). Our results clearly show that CEP-1/P53 does not regulate mitochondrial biogenesis either upon ribosomal stress or increased ambient temperature (Extended Data Fig. 2G-2H).

### SP1 homolog SPTF-3 regulates mitochondrial biogenesis in C. elegans

To unravel the molecular mechanisms connecting the ribosomal stress to mitochondrial biogenesis, we performed two parallel, targeted RNAi screens for TFs regulating mitochondrial biogenesis on-demand upon ribosomal stress or increased temperature. Both screens were carried out using 485 genes annotated as encoding *C. elegans* TFs from different RNAi libraries^20^. The two screens were very conservative and remarkably identified the same transcription factor - SPTF-3, a homolog of mammalian specificity factor 1 (SP1)^21^ (Fig. 3A). While on increased temperature (25°C) we identified another two candidates: *gei-17,* an E3 SUMO-ligase that regulates chromosome segregation ^22^ and W04D2.4, a nematode-specific TF of unknown function (Extended Data Fig. 3A), only SPTF-3 was identified as a suppressor of *mtss-1* expression upon *rps-22* depletion. SPTF-3 is ubiquitously expressed and mainly localizes to the nuclei of somatic tissues (Extended Data Fig. 3B). Other SP1 homologs, SPTF-1 and SPTF-2 failed to reduce *mtss-1* expression upon ribosomal stress (Extended Data Fig. 3C). A previous study, using chromatin immunoprecipitation DNA-sequencing (*ChIP-seq*) showed that SPTF-3 binds to promoter regions of over 100 mitochondrial genes, including around 50% of all OXPHOS subunits, a third of all mitochondrial ribosomal proteins and a large number of other mitochondrial enzymes and factors, including MTSS-1^21^, providing strong validation of our results (Extended Data Table 3). We could also show that *sptf-3* depletion reduced endogenous MTSS-1 protein levels upon ribosomal stress (Fig. 3B). In the absence of SPTF-3, ribosomal stress induced by *rps-22* depletion also failed to increase mtDNA copy number, and steady-state levels of mitochondrial *ctc-1/MT-COX1* transcript (Fig. 3C-3D). Mirroring these results, levels of mitochondrial proteins ATP-1/ATP5A, NUO-1/NDUFV1, and SOD-2/SOD2 increased upon *rps-22* depletion in an SPTF-3-dependent manner (Fig. 3E).

**Figure 3.**
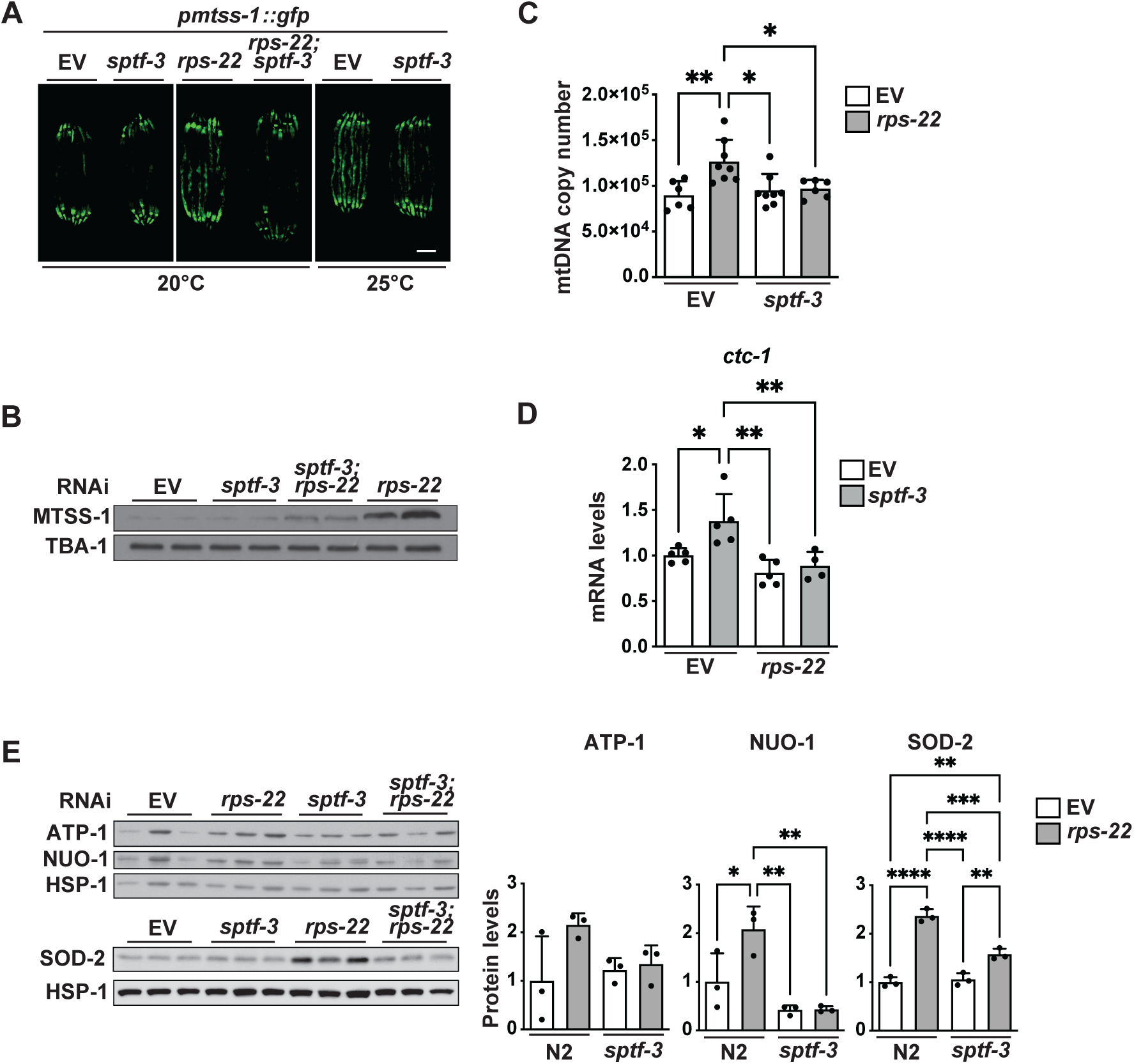
Mitochondrial biogenesis is regulated by SPTF-3. **A)** Fluorescent images of *pmtss-1::gfp* grown on 20°C and 25°C; **B)** Western blot analysis of SPTF-3 steady state levels. TBA-1 was used as a loading control; **C)** mtDNA copy number measured by qPCR at D1; **D)** Steady-state levels of mitochondrial transcript *ctc-1/MT-COX1*; **E)** Steady-state levels of mitochondrial proteins ATP-1/ATP5A, NUO-1/NDUFV1 and SOD-2/SOD2. Western blot (*left*) and quantitative analysis (*right*) are shown. All analyses were performed upon *sptf-3, rps-22* or combined *sptf-3/rps-22* RNAi. Data are presented as mean ± SD. \**p*<0.05, \*\**p*<0.01, \*\*\**p*<0.001, \*\*\*\**p*<0.0001, one-way ANOVA with Tukey post hoc test.

We next developed additional transcriptional reporters for *hmg-5* (*phmg-5::gfp*), a *C. elegans* ortholog of the mammalian mtDNA packaging protein TFAM^23^; and *cts-1* (*pcts-1::gfp*), a homolog of TCA cycle enzyme citrate synthase^24^. In mammals, these two proteins are commonly used to monitor mtDNA levels and mitochondrial mass. On control plates, both reporters showed an increase in GFP fluorescence simply by exposing worms to higher temperatures, which was fully prevented upon *sptf-3* knockdown (Extended Data Fig. 3D-3E). Also, using FACS analysis of a strain expressing a mitochondrially-targeted GFP reporter under the control of gut-specific promoter (*pges-1::mt-gfp*), we could show a small but significant decrease of mitochondrial mass at 25°C upon SPTF-3 depletion (Extended Data Fig. 3F). To exclude the possibility that SPTF-3 directly regulates ribosomal subunits, hence has an indirect effect on mitochondrial biogenesis, we analyzed transcript levels of various genes coding for ribosomal proteins. Indeed, we observed that the absence of SPTF-3 does not affect the expression of ribosomal genes (Extended Data Fig. 3G) suggesting that SPTF-3 has a direct role in regulating mitochondrial biogenesis on-demand.

### SPTF-3 is preferentially translated under stress and regulates several aspects of mitochondrial adaption to stress

Next, we wondered how the activation of SPTF-3 upon ribosomal stress is regulated. SPTF-3 protein levels stabilized upon ribosomal stress caused by *rps-22* RNAi (Fig. 4A), while the transcript levels did not change (Fig. 4B). Depletion of other ribosomal subunits had a similar effect on steady-state levels of SPTF-3 (Extended Data Fig. 4A). As the transcript levels were unchanged, higher steady-state SPTF-3 levels might be a consequence of either decreased turnover or increased protein synthesis. Blocking the general protein degradation pathway by treating worms with proteasome inhibitor bortezomib, did not further increase SPTF-3 levels (Fig. 4C). Remarkably, SPTF-3 protein levels still increased upon *rps-22* knockdown, even in the *ife-2(ok306)* mutant, or in worms treated with cycloheximide, a potent protein synthesis inhibitor (Fig. 4D-4E). This led us to hypothesize that SPTF-3 might be preferentially translated upon stress, similar to ATF4 during the integrated stress response (ISR), although through a different mechanism as phosphorylation of EIF2a was decreased upon *rps-22* depletion (Extended Data Fig. 4B). Mammalian SPTF-3 homolog SP1 accumulates in cells in an internal ribosomal entry site (IRES)-dependent manner^25, 26^ under stress conditions, like during hypoxia or in certain cancer cells. According to IRESite, a database that predicts the existence of IRES sites in transcripts, SPTF-3 is also predicted to possess an IRES sequence^27^. Protein synthesis via initiation at IRESs is favored under conditions when cap-dependent translation is compromised, as seen in many physiological and pathological stress conditions in eukaryotic cells, suggesting that SP1/SPTF-3 might also be regulated in this way.

**Figure 4.**
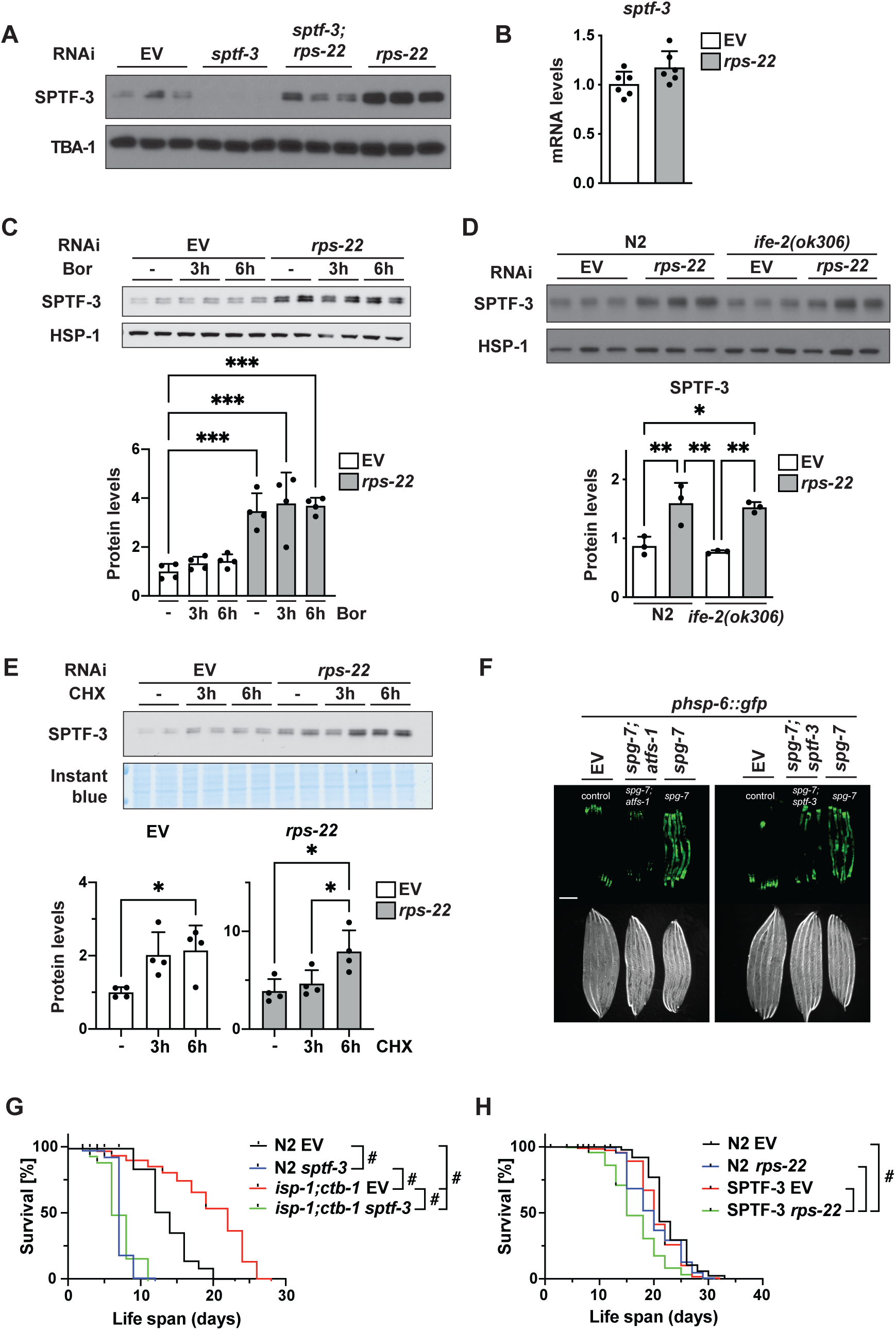
SPTF-3 is translated in a cap-independent manner upon ribosomal stress. **A)** Western blot analysis of SPTF-3 levels upon *sptf-3, rps-22* or combined *sptf-3/rps-22* RNAi; **B)** Steady-state *sptf-3* transcript levels upon *rps-22* RNAi; **C-E)** Steady state levels of STPF-3 measured by western blot (*top*) and quantified using ImageJ analysis (*bottom*) upon: **C)** treatment with proteasome inhibitor bortezomib (Bor); **D)** *rps-22* RNAi in control worms and *ife-2(ok306)* mutant; **E)** treatment with translation inhibitor cycloheximide (CHX) for 3h and 6h. TBA-1 and HSP-1 were used as loading controls; **F)** UPR^mt^ activation measured upon *spg-7/AFG3L2* RNAi using *phsp-6::gfp* reporter, and suppressed either by *atfs-1* or *sptf-3* RNAi; **G)** Lifespans of wild type and *isp-1(qm150)* IV; *ctb-1(qm189)* animals upon *sptf-3* RNAi on 25°C; **H)** Lifespans of wild type and *sptf-3(n4850*) animals upon *rsp-22* RNAi. **A-E)** Data are presented as mean ± SD. \**p*<0.05, \*\**p*<0.01, \*\*\**p*<0.001, **G-H)** ^#^*p*<0.0001. **B, G-H)** unpaired t-test. **C-E)** one-way ANOVA with Tukey post hoc test.

Per SPTF-3 being a stress-regulated TF, its depletion in normal conditions does not induce expression of stress response-markers, e.g., oxidative stress (*gst-4*), endoplasmic reticulum stress (*hsp-4*) or UPR^mt^ - mitochondrial unfolded protein response (*hsp-6*) (Extended Data Fig. 4C). Nevertheless, in the condition of increased mitochondrial stress, caused by depletion of inner mitochondrial m-AAA protease *spg-7/AFG3L2*, SPTF-3 is required for the induction of UPR^mt^ (Fig. 4F and Extended Data Fig. 4D). The suppression of the UPR^mt^ was milder when SPTF-3 was depleted than with the loss of ATFS-1, the direct transcriptional regulator of UPR^mt^ (Fig. 4F), possibly because SPTF-3 indirectly regulates UPR^mt^ by controlling ATFS-1 levels in the cell, as shown by the ChIP-Seq analysis^21^ (Extended Data Table 3). The induction of UPR^mt^ is essential for increased longevity in mitochondrial mutants, such as *isp-1(qm150);ctb-1(qm189)*^28^. We could confirm that loss of *sptf-3* in *isp-1(qm150);ctb-1(qm189)* mutants, strongly reduces ATFS-1 levels and subsequent UPR^mt^ activation, as measured by GFP expression from the *hsp-6/HSPA9* (mitochondrial 70kDa heat shock protein) promoter (Extended Data Fig. 4E-4F). Similarly, *sptf-3* depletion strongly reduces resistance to heat stress in the *isp-1(qm150);ctb-1(qm189)* mutant (Extended Data Fig. 4G).

In agreement with the role in mitochondrial biogenesis and adaptive stress responses, loss of *sptf-3* strongly shortened lifespan in wild-type worms and fully abolished longevity in *isp-1(qm150);ctb-1(qm189)* mutants at 25°C (Fig. 4G). In wild-type worms, SPTF-3 loss had a strong effect on the lifespan at 25°C but not when worms were kept in standard conditions (20°C) (Fig. 4G-4H). Nevertheless, when *sptf-3* was depleted in parallel to *rps-22*, this resulted in a significantly shorter lifespan in worms kept at 20°C (Fig. 4G-4H). Together, these data highlight the role of SPTF-3 as an orchestrator of mitochondria-related adaptive responses upon cellular stress.

### SPTF-3 regulates a conserved starvation response to remodel mitochondria

Mitochondrial function is crucial for maintaining cellular energy homeostasis, and cellular oxidative capacity could be controlled by increasing its mitochondrial content. This is particularly important during times of limited nutrient availability. Although initial studies suggested an increase in mitochondrial biogenesis induced by caloric restriction^10^, most subsequent ones focused on the relationship between nutrient availability and mitochondrial dynamics in different types of cells^29^. We hypothesized that mitochondrial biogenesis is needed when nutrient availability is low, hence both caloric restriction and starvation would induce SPTF-3. Indeed, we could demonstrate that *eat-2(ad1116)* mutants, a genetic model of dietary restriction have elevated SPTF-3 and MTSS-1 protein levels (Fig. 5A). Furthermore, SPTF-3 levels were elevated upon acute starvation of 3 hours, followed by increased *mtss-1* transcription (Fig. 5B-5D). Mirroring this, starvation also induced higher levels of mitochondrial respiratory chain subunits (i.e. ATP-1/ATP5A and NUO-1/NDUFV1) and elevated mitochondrial mass, in a SPTF-3 - dependent manner (Fig. 5E-5F). The mitochondrial biogenesis/remodeling during starvation increases the overall fitness of the animals, as in the absence of SPTF-s, the worms have a significantly shorter lifespan (Fig. 5G). Remarkably, starvation also induced ribosomal stress, as judged by a smaller nucleolar size, suggesting a common mechanism of SPTF-3 induction (Figure 5H).

**Figure 5.**
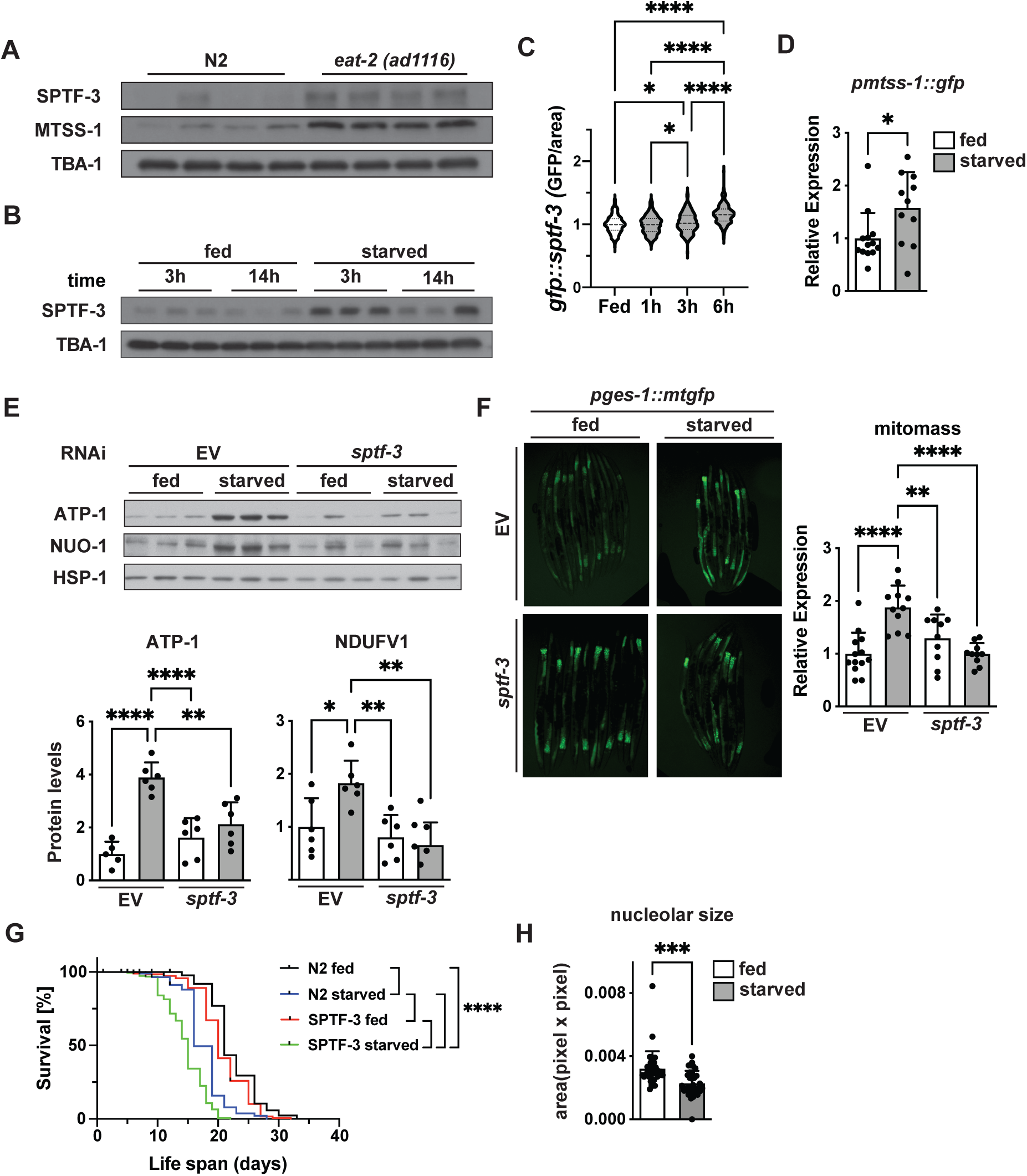
Acute starvation is inducing similar SPTF-3 response. **A)** Western blot analysis of SPTF-3 and MTSS-1 levels in *eat-2(ad1116)* mutants; **B)** Western blot analysis of SPTF-3 upon acute (3h) and chronic (14h) starvation; **A-B)** TBA-1 was used as loading control; **C)** Analysis of SPTF-3 levels using reporter strain expressing *psptf-3::gfp::sptf-3 upon* 1h, 3h and 6h starvation; **D)** *mtss-1* expression levels measured using *pmtss-1::gfp* reporter in upon 3h starvation; **E)** Steady-state levels of mitochondrial proteins ATP-1/ATP5A and NUO-1/NDUFV1 upon 3h starvation in the presence of *sptf-3* RNAi. HSP-1 is used as a loading control. Western blot (*top*) and quantitative analysis (*bottom*) are shown; **F)** Total mitomass measured using a *pges-1::mtgfp* reporter upon *sptf-3* RNAi and 3h starvation. Representative florescent images (*left*) and quantitative analysis (*right*); **G)** Lifespans of wild type and *sptf-3(n4850)* mutant upon starvation during D1 of adulthood; **H)** Nucleolar size measured by the expression of the GFP-tagged nucleolar protein FIB-1 using *pfib-1::fib-1::gfp* reporter strain. Data are presented as mean ± SD. \**p*<0.05, \*\**p*<0.01, \*\*\**p*<0.001, \*\*\*\**p*<0.0001. One-way ANOVA with Tukey post hoc test was used in C, E and F. Unpaired t-test was used in D, G and H.

Importantly, autophagy does not play a role in this pathway, as SPTF-3 depletion did not affect either mitophagy or general autophagy upon starvation (Extended Data Fig. 5A-5B). We could show that loss of SPTF-3 did not increase mitophagy in starved animals, as measured by a dual fluorescent pH-sensitive biosensor with a mitochondrial tag that, upon mitophagy induction, mainly quenches the green fluorescence, thus reducing the GFP to dsRed ratio (Extended Data Fig. 5A). Similarly, no change in autophagy flux was detected regardless if SPTF-3 was present in starved worms, using antibodies raised against membrane-bound LGG-1/LC3 or cytoplasmic phosphatidylethanolamine (PE)-conjugated PE-LGG-1/LC3, as well as release of soluble GFP (Figure S5B).

Next, we asked whether a similar function is conserved for SP1, a mammalian homolog of SPTF-3. It was previously shown that SP1 acts as a transcriptional regulator of many genes encoding OXPHOS proteins, including most *COX* subunits and *TFAM* by binding their promoters^30^ and is necessary for starvation adaption response by regulating aspartate catabolism genes^31^. Here, we showed that SP1 levels increase upon glucose and amino acid starvation in HeLa cells (Figure 6A). Following the increase in SP1, the mtDNA levels were also induced by both interventions leading to nutrient deprivation (Figure 6B). This was mirrored by an increase in the mtDNA packaging protein TFAM (Figure 6C). Importantly, the levels of PGC-1a, the transcriptional regulator of mitochondrial biogenesis on-demand were not changed, suggesting that SP1 and not PGC-1α play a role in this process (Figure 6C).

**Figure 6.**
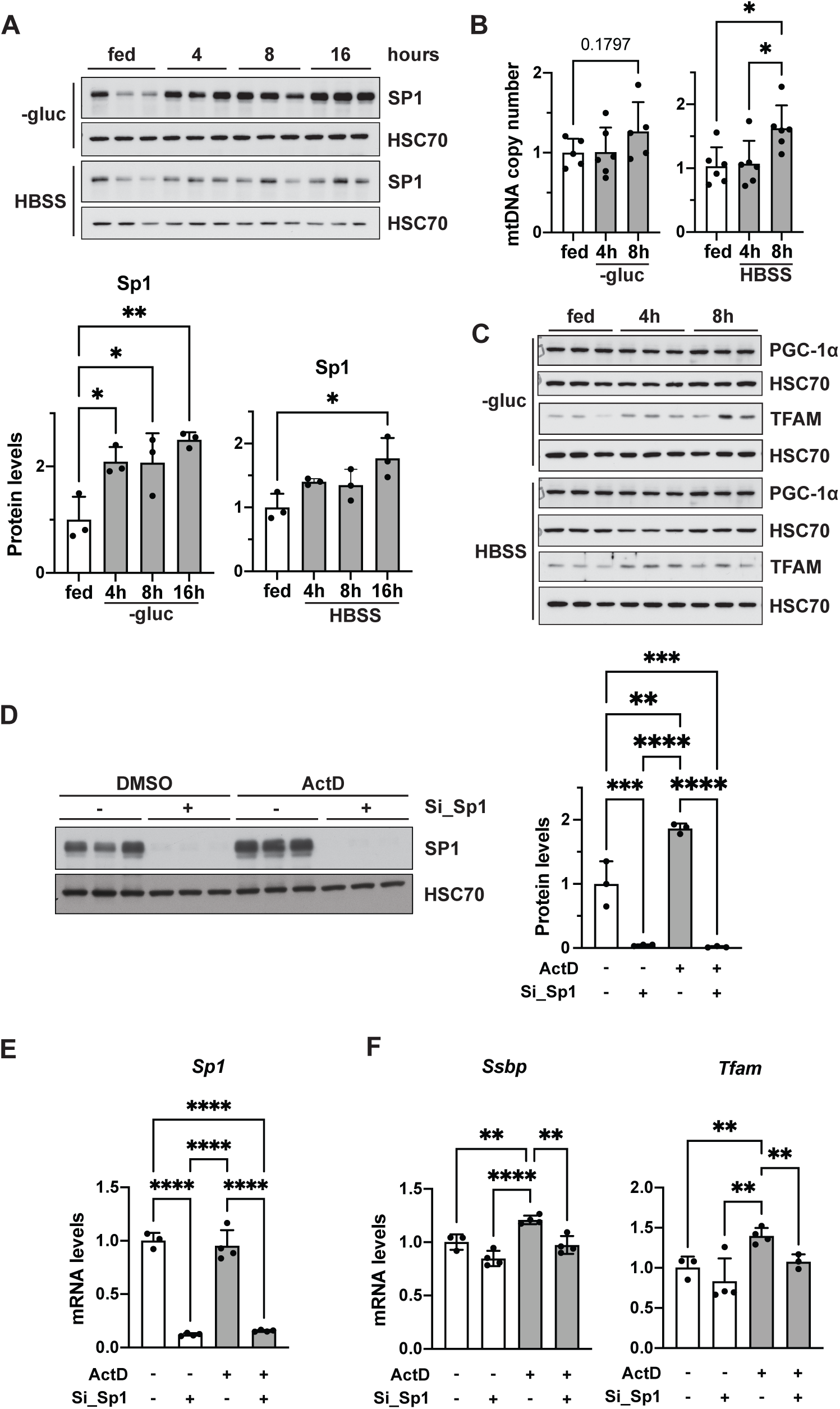
STPF-3 stress-adaptive response is conserved in mammals. A-C) Experiments were performed in HeLa cells upon glucose (-gluc) or amino acid (HBSS) starvation, for times indicated in panels: **A)** Steady state levels of SP1 upon nutrient starvation for 4h, 8h and 16h. HSC70 was used as a loading control; **B)** mtDNA copy number measured upon nutrient starvation for 4h, 8h; **C)** Western blot of TFAM and PGC-1α in cells starved for 4h and 8h; **D-F)** Experiments were performed in HeLa cells upon treatment with Actinomycin D (ActD) for 24h in the presence of *Sp1* siRNA: **D)** Steady-state levels of SP1; **E)** Steady-state SP1 transcript levels; **F)** Steady-state transcript levels of mitochondrial *Ssbp* and *Tfam*. Data are presented as mean ± SD. \**p*<0.05, \*\**p*<0.01, \*\*\**p*<0.001,\*\*\*\**p*<0.0001, one-way ANOVA with Tukey post hoc test.

Echoing the results in *C. elegans*, treatment with ActD that induces ribosomal stress also resulted in the upregulation of SP1 levels, but not its transcripts (Figure 6D – E, respectively). Moreover, the mitochondrial biogenesis response also seems to be conserved in these conditions as we found that transcript levels of mammalian homologs of *mtss-1/Ssbp* and *hmg-5/Tfam* are also upregulated by ActD treatment in an SP1-dependent manner (Figure 6F). Thus, the SPTF-3-mediated stress-adaptive response leading to increased mitochondrial biogenesis is conserved in mammals through the action of its homolog SP1.

## Discussion

Mitochondrial plasticity and the capacity to rapidly respond to increasing energy needs rely heavily on mitochondrial biogenesis on-demand. Whereas much progress has been achieved in understanding the transcriptional regulation of mitochondrial genes in mammals, little is known about the transcriptional regulation of mitochondrial biogenesis on-demand in invertebrates, particularly in the absence of pivotal TFs (e.g., PGC-1a and NRF1-2) identified in mammals^11^. Here, we show that mitochondrial biogenesis is induced by disturbances in ribosomal assembly, which leads to skewed cytoplasmic translation resulting in the preferential synthesis of stress-induced transcription factor SPTF-3. Although ribosomal biogenesis stress also causes P53 activation upon accumulation of unassembled ribosomal proteins, we showed that CEP-1/P53 is dispensable for an increase in mitochondrial biogenesis. By analyzing data from a previous study^21^ we could show that SPTF-3 directly binds promoters of mitochondrial genes, in particular the ones encoding OXPHOS subunits and proteins involved in mitochondrial gene expression. Here we demonstrated that SPTF-3 regulates levels of multiple mitochondrial proteins in the stress conditions like increased temperature, caloric restriction, or acute starvation, all of which induce an adaptive increase of mitochondrial mass, thus rising organismal fitness and survival.

SPTF-3 is a member of the Specificity Protein/Krüppel-like factor (SP/KLF) family of C2H2-type zinc-finger transcription factors^32^. In *C. elegans* this family consists of six TFs (SPTF-1 to 3 and KLF-1 to 3), while mammals have nine SP and 16 KLF subfamily members^32^. All of them play key roles in multiple biological processes including stem cell maintenance, cell proliferation, embryonic development, tissue differentiation, and various aspects of metabolism, hence it is not surprising that their dysregulation has been implicated in several human diseases^32^. We recently described the role of another transcription factor from the SP/KLF family as the major regulator of adaptive stress response in *C. elegans*. We identified KLF-1 as a main mediator of a cytoprotective response that dictates longevity induced by reduced mitochondrial function^33^. We could demonstrate that KLF-1 activates genes involved in the xenobiotic detoxification program and identified cytochrome P450 oxidases as longevity-assurance factors of mitochondrial mutants^33^. Our further results showed that KLF-1 translocation to the nucleus and the activation of the signaling cascade is dependent on the mitochondria-derived hydrogen peroxide (H_2_O_2_) produced during late developmental phases where aerobic respiration and somatic mitochondrial biogenesis peak^34^. Curiously, we now identified SPTF-3, a second transcription factor from the same family to regulate mitochondrial adaptive stress response, suggesting evolutionary conservation and possible divergence from a common ancestral TF.

Nematode SPTF-3 has not been extensively studied, although it was demonstrated that its loss causes defects in the development of the egg-laying system, oocyte production, and embryonic morphogenesis^35^. SPTF-3 further controls the programmed cell death of several motor and sensory neurons by transcriptionally activating both caspase-dependent and -independent apoptotic pathways^21^. The transcription complex formed by SPTF-3 mammalian homolog SP1 is highly variable and contains multiple other transcription factors and co-factors, hence can activate different genes in a cell- and context-dependent manner^32^. Remarkably, SP1 is one of the first transcription factors shown to be implicated in the regulation of gene expression of nDNA-encoded mitochondrial proteins^36^. SP1 recognition sites can be found within the promoter region of TFAM, TFB1M, TFB2M, and POLG2, factors essential for mtDNA replication/transcription^37^. SP1 was also shown to induce the expression of all 13 cytochrome c oxidase (COX) subunits^30^. Consistently, genes of many OXPHOS subunits, including factors involved in their expression and some TCA enzymes were also identified as targets of the nematode’ SPTF-3 in ChIP-Seq experiments^21^. We could demonstrate a tight regulation of *mtss-1* and *cts-1* expression by SPTF-3, suggesting that SP1 is indeed its evolutionarily-conserved functional homolog.

Remarkably, the changes induced by increased temperature or ribosomal biogenesis stress on overall mitochondrial mass were always milder than the upregulation of components involved in mitochondrial gene expression (*mtss-1*, *hmg-5*) or OXPHOS subunits (ATP-1 and NUO-1). Although we referenced this change as “mitochondrial biogenesis”, the more precise term for what occurs to the mitochondrial network might be “mitochondrial or OXPHOS remodeling” as the majority of mitochondrial genes regulated by SPTF-3 are either OXPHOS subunits or directly involved in OXPHOS biogenesis^21^. Therefore, we propose that SPTF-3 might have a specific role in the OXPHOS remodeling on-demand, thus galvanizing the overall increase in mitochondrial biogenesis to meet the new (patho-)physiological conditions. Similarly, the SP1 role in metabolic remodeling has been previously recognized for most tumors, where overexpression of SP1 (also SP2, SP3) is a negative prognostic factor for survival in patients^38^.

*Our results demonstrated that increased cellular stress caused by inhibition of ribosomal biogenesis enhances SPTF-3 translation, most probably through an IRES-mediated mechanism. A similar pathway was described for SP1 regulation under ischemia-induced H_2_O_2_ stress*^25^ *and during tumorigenesis*^26^*. Under normal growth conditions, IRES-mediated translation utilizes the common ternary complex-dependent translation initiation, whereas during stress IRES-containing RNAs are translated by an EIF5B-dependent mechanism*^39^. Defective ribosomal biogenesis also causes a decrease in CAP-dependent translation, paralleled by an increase in translation of IRES-containing transcripts, or transcripts with shorter and less structured 5’-UTRs^39^. Curiously, in *Drosophila melanogaster*, caloric restriction was shown to reduce the global translation rate, while enhancing the translation of many mitochondrial transcripts as they possess less structured 5’-UTR^40^. Consequently, the reduction of OXPHOS subunits diminished the lifespan extension effect of CR in fruitflies^40^. Nutrient stress requires adaptation through transcriptional responses that affect organismal growth and survival. Caloric restriction in the yeast prolongs lifespan by shifting carbon metabolism towards OXPHOS, thus significantly increasing respiration rate^41^. Similarly, increased metabolic rate, dependent on functional OXPHOS, is essential for the lifespan extension in *C. elegans* grown in reduced food conditions^42^. This response is conserved across evolution as CR in rats increases the number of mitochondria in the liver^43^. We could show that SPTF-3 is upregulated in response to CR and acute starvation where it regulates adaptive mitochondrial biogenesis response. In parallel, SPTF-3 also directly regulates the expression of *atfs-1,* the major transcriptional switch of UPR^mt^ that controls mitochondrial proteostasis^44^. In this way, SPTF-3 shields mitochondrial fitness by regulating both mitochondrial biogenesis and response to photostatic stress. This increases fitness during starvation, ensures longevity in *isp-1(qm150);ctb-1(qm189)* mutants, and allows animals to survive high ambient temperatures. Other studies also propose alterations of cytosolic translation as a link coupling nutrient deprivation, mitochondrial biogenesis, and longevity^45–48^, possibly because translational regulation of selective mRNAs is a sophisticated way to quickly adjust protein concentration in response to stress stimuli. The pivotal role in the regulation of translation events is often attributed to the nutrient-sensing serine/threonine protein kinase TOR (target of rapamycin)^49, 50^. Indeed, it was shown that mTORC1 promotes the selective translation of mitochondrial genes by phosphorylating 4E-BP^51^ and mediates the formation of the translation preinitiation complex via S6 kinase (S6K)^52^. Nevertheless, we could demonstrate that mTORC1 and mTORC2 complexes do not play a major role in the regulation of SPTF-3-mediated induction of mitochondrial biogenesis upon increased temperature or ribosomal stress. Taken together, we identified a novel mechanism of regulation of mitochondrial biogenesis on-demand and put SPTF-3 in the middle of this pathway critical for many stress conditions that require remodeling of the mitochondrial network. We established that SPTF-3 also regulates mitochondrial proteostasis by controlling UPR^mt^ transcriptional activation. As a result, SPTF-3 safeguards mitochondrial well-being by regulating two parallel programs - mitochondrial biogenesis and UPR^mt^. This is particularly important in the early steps of adaptation to nutrient deprivation when steady mitochondrial function ensures optimal energy production. Remarkably, we could demonstrate an evolutionarily conserved role of SP1, the mammalian homolog of SPTF-3, in the regulation of mitochondrial biogenesis response to acute nutrient starvation. Therefore, we propose that SP1 and other members of the SP/KLF family of transcription factors might play a much bigger role in the regulation of mammalian mitochondrial adaptive responses to various stresses than assumed so far, hence should be considered as putative therapeutic targets.

## Acknowledgments

The authors wish to thank H. Robert Horvitz (Massachusetts Institute of Technology, Cambridge, USA) for providing us with the anti-SPTF-3 (M82) antibody and Nils-Göran Larsson for providing the anti-TFAM antibody.

The work was supported by Aleksandra Trifunovic’s grants from the German Research Foundation (Deutsche Forschungsgemeinschaft - DFG) - SFB 1218 Projektnummer 269925409 and Centre for Molecular Medicine Cologne, University of Cologne (C15).

## Material & Methods

### Maintenance and strains of *C. elegans*

Populations of *C. elegans* were cultured on *E.coli* OP50 bacteria-seeded NGM plates, according to standard protocols unless otherwise stated ^53^. The following strains were used in this study: Bristol N2 (wild type, N2), ATR1010; *isp-1(qm150)* IV; *ctb-1(qm189),* zcls13 [*hsp-6p::gfp* V], ATR4094; unc-119;Ex(*pmyo-3::tomm-20-Rosella; Cb unc- 119*), CL2166; dvls19[(*pAF15)gst-4p::gfp::nls*] III, DA1116; *eat-2(ad1116)* II, KX15; *ife-2(ok306)*, MAH235; sqls19 [*hlh-30p::hlh-30::gfp, rol-6(su1006)*], MQ989; *isp- 1(qm150)* IV; *ctb-1(qm189),* MT19372; *sptf-3(n4850)* I; nls283*[gcy-10p::GFP + lin- 15(+)]*, MT19859; nls431[*gfp::sptf-3*] X, RB967; Y81G3A.3*(ok871)* II, SJ4005; zcls4 [*hsp-4p::gfp*] V, SJ4058; zcls9 [*hsp-60p::gfp*] V, SJ4100; zcls13 [*hsp-6p::gfp* V] and SJ4143; zcls14[*ges-1::gfp*(mit)], VB0633; *rsks-1(sv31)* III. ATR0026; Ex(*pcts-1::gfp)*, ATR0027; Ex(*phmg-5::gfp),* ATR2636; Ex(*pmtss-1::gfp*), and ATR4057; Ex(*pfib-1::fib- 1::gfp*) were generated by injecting *pcts-1::gfp, phmg-5::gfp, pmtss-1::gfp* and *pfib- 1::fib-1::gfp* (40 ng/µl), respectively, *pGH8* (20 ng/µl) and *pRF4* (40 ng/µl) into N2 animals. Plasmid mixtures were injected using standard procedures ^54^. ATR2636 was subsequently integrated by X-ray and outcrossed to create ATR0011. ATR0011 was crossed into KX15 to create ATR0022. ATR0011 was crossed into RB967 to create ATR0024. ATR0011 was crossed into VB0633 to create ATR4107.

### RNAi knockdown

RNAi knockdown was performed as described previously ^55^. The RNAi clones used in this study were obtained from the Ahringer ^55^ and Vidal RNAi library ^56^ and are listed in Table 1. Bacteria were grown in Luria broth medium to OD_595_ = 0.5, then 1 mM IPTG was added, and bacteria were induced for 3 h at 37 °C and consequently seeded on NGM plates containing 100 µg/ml ampicillin, 20 µg/ml tetracycline and 1 mM IPTG. Animals were treated with RNAi from hatching. For control, bacteria carrying the empty vector (EV) L4440 were used.

### Lifespan analysis

Worms were grown at 20°C or 25°C and exposed to RNAi from hatching unless otherwise stated. D1 was defined as day 1 of the lifespan. On day 1, 25 animals were transferred to each NGM plate, for a total of 100 worms per condition. Acute starvation was performed by transferring D1 worms to plates without bacteria for 30h. Worms that escaped the plate or died due to protrusion or internal hatching were censored from the experiment. Compilation of lifespan data and statistical analysis are enlisted in Extended Data Table 4.

### Heat stress assay

D1 worms were exposed to 37°C for 6h in a pre-heated water bath. Then the worms were allowed to recover at 20°C on a NGM agar plate with bacteria and after 1h scored by survival by prodding gently. 30 worms per experiment were scored per condition.

### Cell culture experiments

HeLa cells were cultured in DMEM medium containing 4 mM L-Glutamine, 1 mM sodium pyruvate, 100 units/mL penicillin-streptomycin and 10% fetal bovine serum (FBS). For the starvation experiments, cells were cultivated for 24h at 37 °C in culture medium, washed with PBS, and either DMEM medium without glucose or HBSS medium was used. For the gene silencing of *Sp1* and negative control, 1.2 × 10^5^ HeLa cells were transfected in a reverse transfection protocol with Lipofectamine RNAiMAX (Thermo Fisher Scientific) and *Sp1* siRNA (purchased from Eurogentec) at a dose of 1 nM in 12 wells following the manufacturer’s instructions. siRNA sequence for *Sp1* was CCAACAGAUUAUCACAAAU. After 24h cultivation at 37 °C in culture medium, actinomycin D (Sigma Aldrich) was added to a final concentration of 50 nM and incubated for additional 24 h. After the incubation, the cells were collected and used for RNA or protein isolation.

### RNA isolation and qPCR

Animals or cells were collected, respectively, from one full NGM agar plate or from one well from 6 well plates and total RNA was isolated with Trizol (Invitrogen). DNAse treatment was achieved using DNA-free^TM^, DNAse treatment and removal kit (Ambion, Life Technologies) or DNAseI (NEB), according to the manufacturer’s protocol. Amount of total RNA was quantified by spectrophotometry and 0.8 µg total RNA was reversely transcribed using the High-Capacity cDNA Reverse Transcription Kit (Applied Biosystems), with the following PCR conditions: 95°C for 3 min, followed by 40 cycles of 95°C for 5 sec, 60°C for 15 sec. Detection of amplified products was performed using Brilliant III Ultra Fast SYBR Green qPCR Master Mix (Agillent Technologies) or PowerUp Syber Green master mix (Thermo Fisher) and normalization was against Y45F10D.4. Used primers are enlisted in Table 2. Data was represented using △△Ct and for each condition, five independent replicates were used for worms, 4 -5 independent replicates were used for cells.

### Western blotting

Protein sample preparations from worms were performed by collecting worms from three full plates and washing them at least three times in M9 buffer. Worm pellet was frozen in liquid nitrogen and, depending on the amount of worm pellet, liquefied in 100- 200 µL worm lysis buffer (25 mM Tris-HCL pH 7.4, 0.15 M NaCl, 1 mM EDTA, 1% NP- 40, 0.5% SDS, 10 mM DTT and proteinase inhibitor cocktail (Roche)) or RIPA buffer with proteinase inhibitor cocktail (Roche). PhosStop (Roche) was added when phospho-antibody was used. Cells were collected by trypsinization and washed in PBS. Cells were frozen in liquid nitrogen. Samples were sonicated using the Bioruptor^R^ Sonication device (Diagenode) six times at 30/60 cycles. Samples were then centrifugated for 15 min at 16,000 *x g* at 4 °C and the supernatant was used. Protein concentration was measured with Bradford assay. Western blotting was performed using antibodies, with 1:2000 dilution unless otherwise stated, against ATP-1 (ATP5A, Abcam, #ab14748), GFP (OriGene, #TP401), HSP-1 (HSC70 (B6), 1:4000, Santa Cruz, #sc-7298), HSP-6 (Grp70, Abcam, #82591), HSP-60 (ATCC), mnSOD (Merck, 06-984), MTSS-1 (SSBP1, Atlas Antibody, #HPA002866), NUO-1 (NDUFV1, Proteintech, #11238-1-AP), PGC-1α (Santa Cruz, #sc-13067), Phospho-eIF-2α (eIF2α, Abcam, #ab32157), Phospho-(Ser/Thr) Phe (Cell Signaling, #9631), Puromycin (Merck, #MABE343), SP1 (Abcam, #ab124804), SPTF-3 (M82) (gift from H.R. Horvitz), TBA-1 (Tubulin, Sigma Aldrich, #6074) and TFAM (gift from Nils-Göran Larsson). Data were normalized to either TBA-1 or HSP-1.

### Ribosomal profiling

Previously used method was optimized to resolve the polysomes and ribosomal subunits^48^. 100 µl of gently pelleted D2 worms were homogenized on ice in 300 µL of solubilization buffer (300 mM NaCl, 50 mM Tris-HCl (pH 8.0), 10 mM MgCl_2_, 1 mM EGTA, 200 µg heparin per mL, 400 U RNAsin per mL, 1.0 mM phenylmethylsulphonyl fluoride, 0.2 mg cycloheximide per mL, 1% Triton X- 100, 0.1% Sodium deoxycholate) by 60 strokes with a Teflon homogenizer. 700 µL additional solubilization buffer was added, vortexed briefly, and placed back on ice for 10 min before centrifuging the sample at 20 000 *g* for 15 min at 4 °C. Approximately 0.9 mL of the supernatant was applied to the top of a 10–50% sucrose gradient in high salt resolving buffer (140 mM NaCl, 25 mM Tris-HCl (pH 8.0), 10 mM MgCl_2_) and centrifuged in a Beckman SW41Ti rotor (Beckman Coulter, Fullerton, CA, USA) at 180,000 *g* for 90 min at 4 °C. Gradients were fractionated with continuous monitoring of absorbance at 260nm.

### Oxygen consumption rates

Oxygen consumption rates were measured using the Oxygraph-2k high resolution respirometry system (Oroboros Instruments GmbH). Per condition 300 animals per condition were manually transferred into the Oxygraph chamber and oxygen consumption was analyzed for at least 15 min. Experiment was performed at least six times and conditions were alternating between the chambers. The slopes of the linear portions of the plots were used for calculation and data were analyzed using DatLab4 software (version 4.3).

### BioSorter

GFP intensity in large populations of worms was analyzed by BioSorter. D1 animals were collect with M9 buffer and washed multiple times before using the BioSorter®INSTRUMENT. FlowPilot™ software was used for analysis and at least 300 animals were used per condition.

### Microscopy

Worms were paralyzed with 50 mM sodium azide and mounted on 2% agarose pads. Images were captured using an Axiolmager Z.1 epifluorescence microscope with Zen (blue edition) software and analyzed using Image J (National Institutes of Health).

### Statistical analysis

To analyze statistical significance, a two-tailed unpaired Student’s t-test, Chi-square test, or one-way ANOVA with Tukey post hoc test were used, unless otherwise stated. Error bars are represented as the standard deviation (SD) and significance is stated by *p*<0.05 and represented by stars (\**p*<0.05, \*\**p*<0.01, \*\*\**p*<0.001, \*\*\*\**p*<0.0001) or symbol (^#^*p*<0.0001). Unless otherwise stated, all experiments were performed three times independently of each other.

## Notes

### Competing Interest Statement

The authors have declared no competing interest.

