## Supplementary files for "SPTF-3/SP1 orchestrates mitochondrial biogenesis upon ribosomal stress and acute starvation"

#### Extended Data Figures and Tables

#### Extended Data Figures 1 - 5

#### Extended Data Tables 1 - 4

#### Extended Data Fig.1

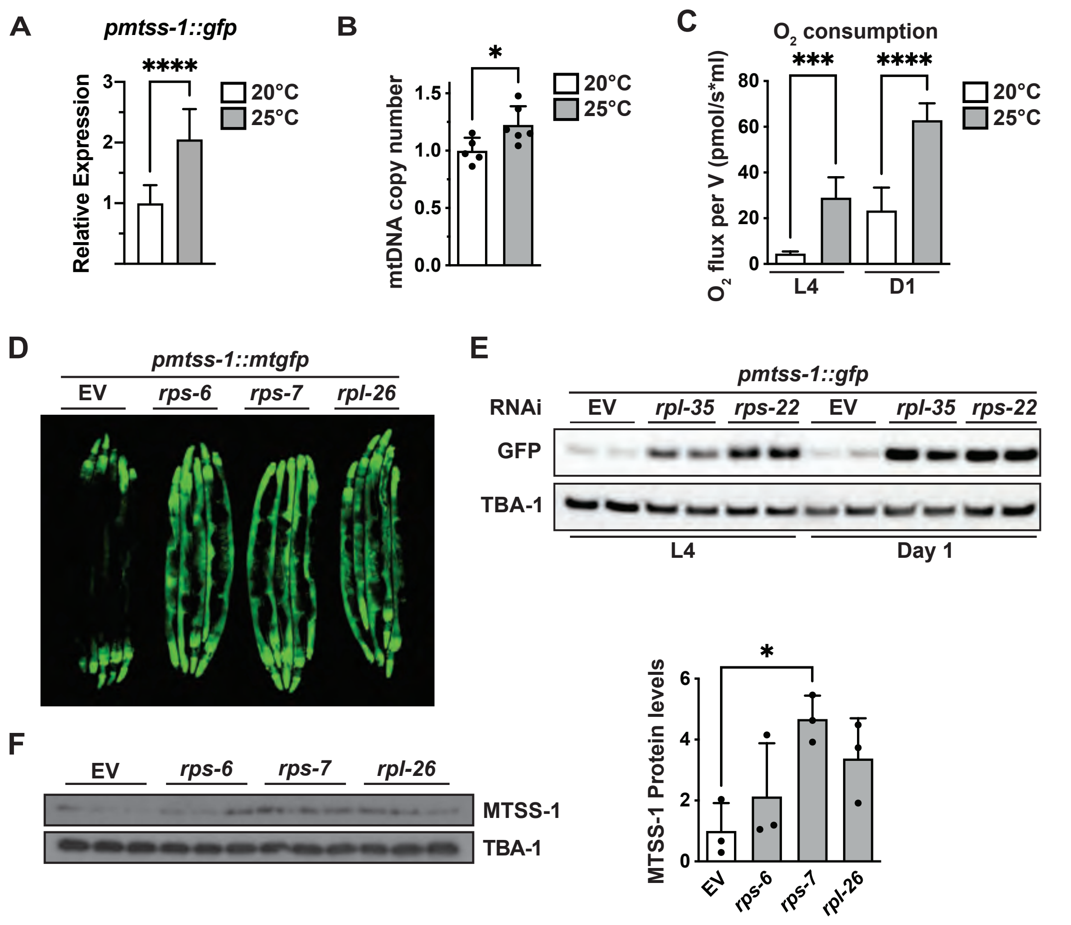


**Extended Data Fig.1 Ribosomal-dependent MTSS-1 level is a readout for mitochondrial biogenesis.**

**A-C)** Animals were grown on either 20°C or 25°C: **A)** Levels of *pmtss-1::gfp* analysed by BioSorter®; **B)** mtDNA copy number measured by qPCR; **C)** Basal oxygen consumption analysed by Oroboros in L4 and D1 animals; **D)** *mtss-1* expression levels using *pmtss-1::gfp* reporter upon *rps-6, rps-7* or *rpl-26* RNAi; **E)** Western blot with steady-stated levels of GFP in animals expressing *pmtss-1::gfp* upon *rpl-35* and *rps-22* RNAi; **F)** Steady-stated levels of MTSS-1 upon *rps-6, rps-7* and *rpl-26* RNAi. Western blot (*left*) and quantitative analysis (*right*) are shown; **E-F)** Loading control was TBA-1. Data are presented as mean ± SD. $^{*}p<0.05$, $^{**}p<0.01$, $^{***}p<0.001$, $^{****}p<0.0001$. **A-B)** unpaired t-test, **C, F)** one-way ANOVA with Tukey post hoc test.

### Extended Data Fig.2

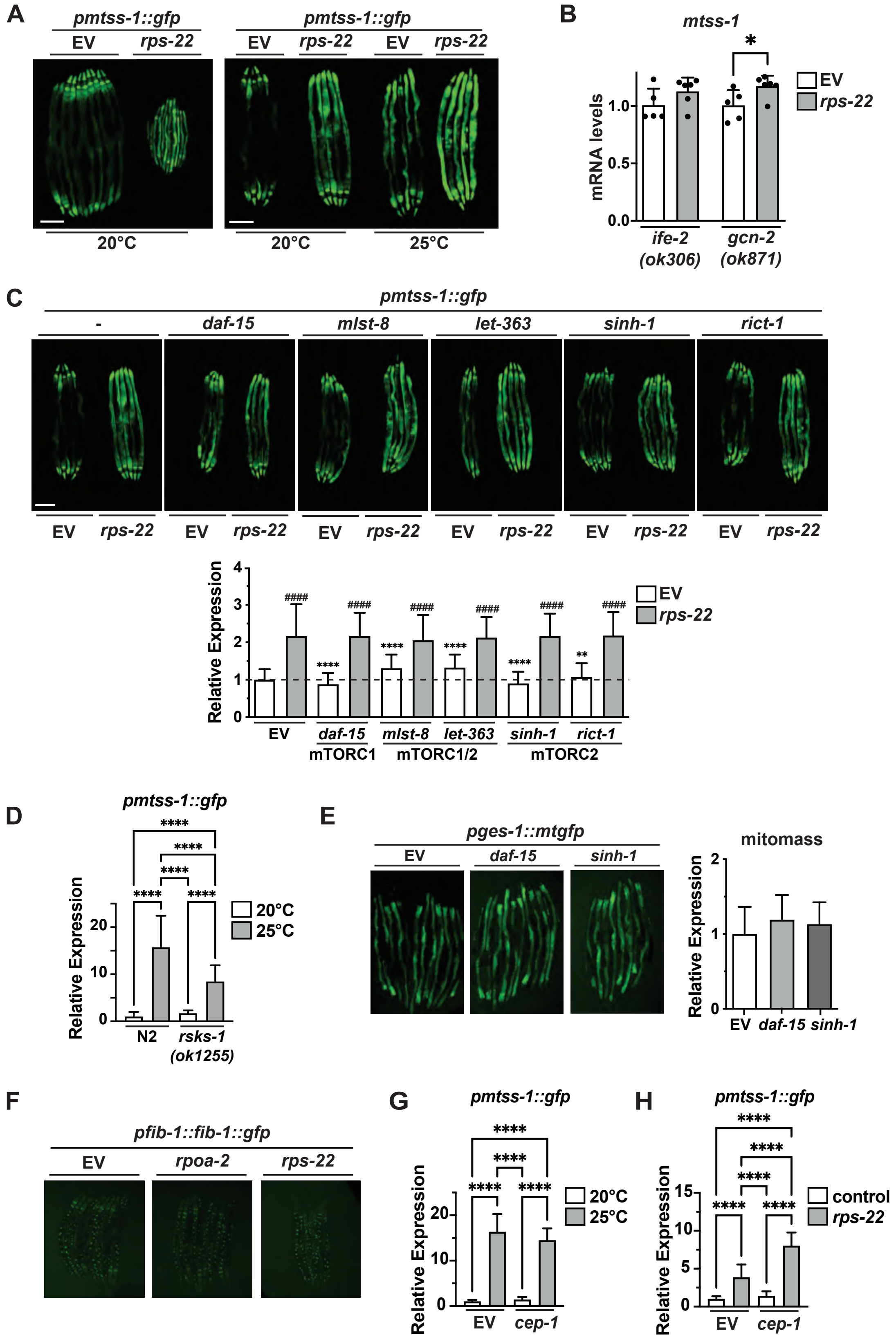

**Extended Data Fig.2 Upregulation of MTSS-1 is translation and mTORC-independent.**

**A)** Fluorescent images of *pmtss-1::gfp* reporter strain upon *rps-22* RNAi. Animals treated with undiluted *rps-22* RNAi on 20°C (*left*) or with diluted 1:1 *rps-22*/L4440 RNAi on either 20°C or 25°C (*right*) are shown; **B)** Steady-state *mtss-1/SSPB1* transcript levels in *gcn-2(ok871)* and *ife-2(ok306)* upon *rps-22* RNAi; **C)** *mtss-1* expression levels measured using *pmtss-1::gfp* reporter upon RNAi against different mTORC1, mTORC1/mTORC2 and mTORC2 subunits and *rps-22* RNAi (*up*) and quantified using BioSorter® (*bottom*); **D)** Quantification of *pmtss-1::gfp* reporter strain in *rsks-1(ok1255)* mutant on either 20°C or 25°C. **E)** Fluorescent images of *pges-1::mtgfp* reporter strain upon *daf-15* or *sinh-1* RNAi (*left*) and quantitative analysis (*right*); **F)** Representative images of nucleolar size of *pfig-1::fib-1::gfp* reporter upon *rpoa-2* RNAi. Quantified in Figure 2H; **G-H)** The expression levels of *mtss-1/SSBP-1* by using *pmtss-1::gfp* reporter strain upon *cep-1* RNAi: **G)** Animals on 20°C and 25°C; **H)** Animals upon *rps-22* RNAi. **B, D-E, G-H)** Data are presented as mean ± SD. $^{*}p<0.05$, $^{**}p<0.01$, $^{***}p<0.001$, $^{****}p<0.0001$. **C)** * are representing significance in comparison with control EV, # are representing significance in comparison between EV and *rps-22* RNAi. **B-C)** unpaired t-test. **D-E, G-H)** one-way ANOVA with Tukey post hoc test.

### Extended Data Fig.3

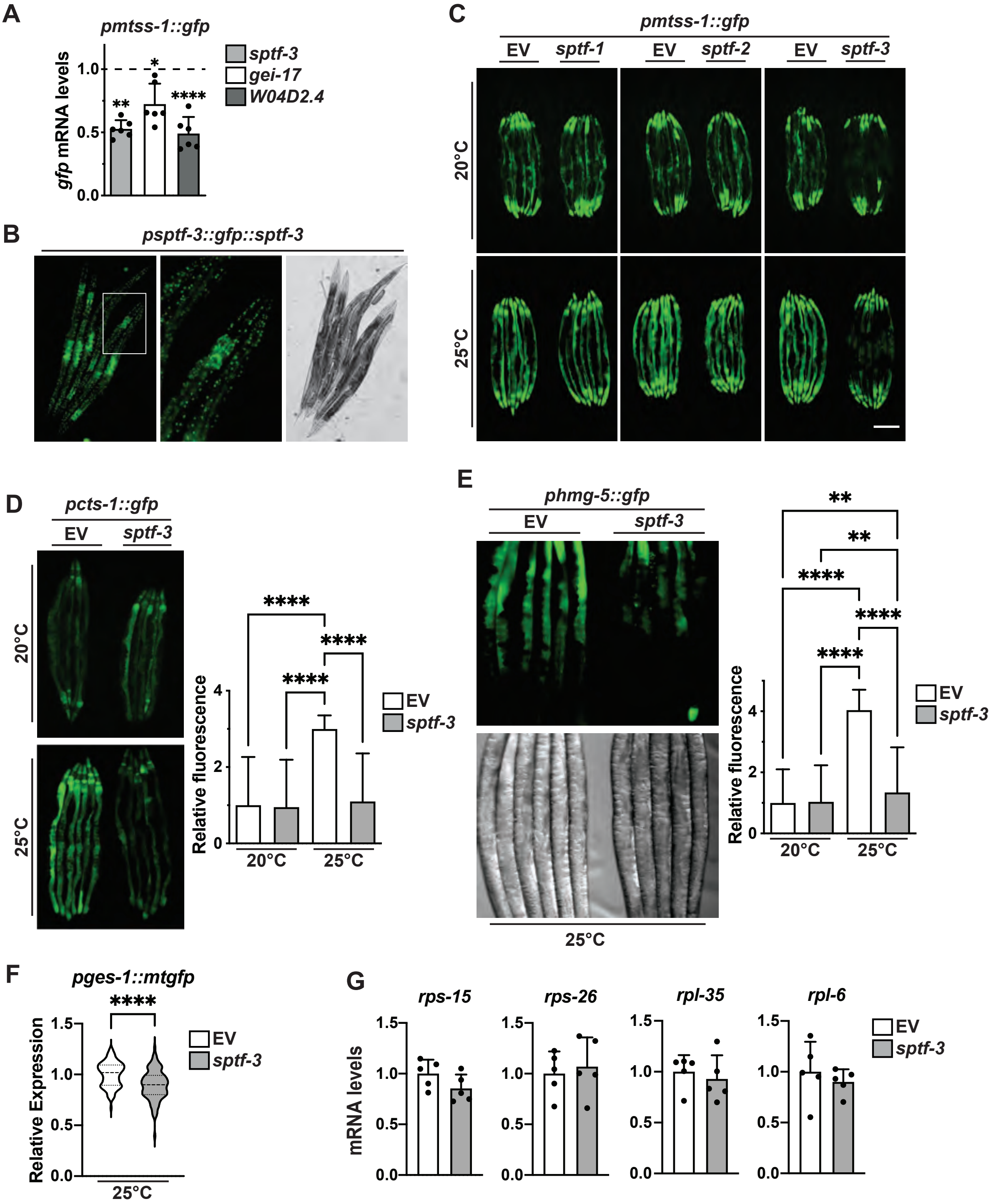

**Extended Data Fig.3 SPTF-3 specific is orchestrating mitochondrial biogenesis.** **A)** Steady-state *gfp* expression levels in *pmtss-1::gfp* reporter strain upon *sptf-3, gei-17,* or *W04D2.4* RNAi measured by qPCR; **B)** Representative florescent and DIC images of *psptf-3::gfp::sptf-3* expression pattern; **C)** Fluorescent images of *pmtss-1::gfp* reporter strain upon *sptf-1, sptf-2* or *sptf-3* RNAi on either 20°C or 25°C; **D-E)** Representative florescent images (*left*) and quantitative analysis by BioSorter® (*right*) of animals upon *sptf-3* on either 20°C or 25°C: **D)** *cts-1* expression levels measured using *pcts-1::gfp* reporter strain; **E)** *hmg-5* expression levels measured using *phmg-5::gfp* reporter strain; **F)** Total mitomass analysed using *pges-1::mtgfp* upon *sptf-3* RNAi on 25°C; **G)** Steady-state transcripts levels of ribosomal subunit *rps-15, rps-26, rpl-35* and *rpl-6* upon *sptf-3* RNAi. Data are presented as mean ± SD. $^{*}p<0.05$, $^{**}p<0.01$, $^{***}p<0.001$, $^{****}p<0.0001$. **A, D-E)** one-way ANOVA with Tukey post hoc test. **F-G)** unpaired t-test,

#### Extended Data Fig.4

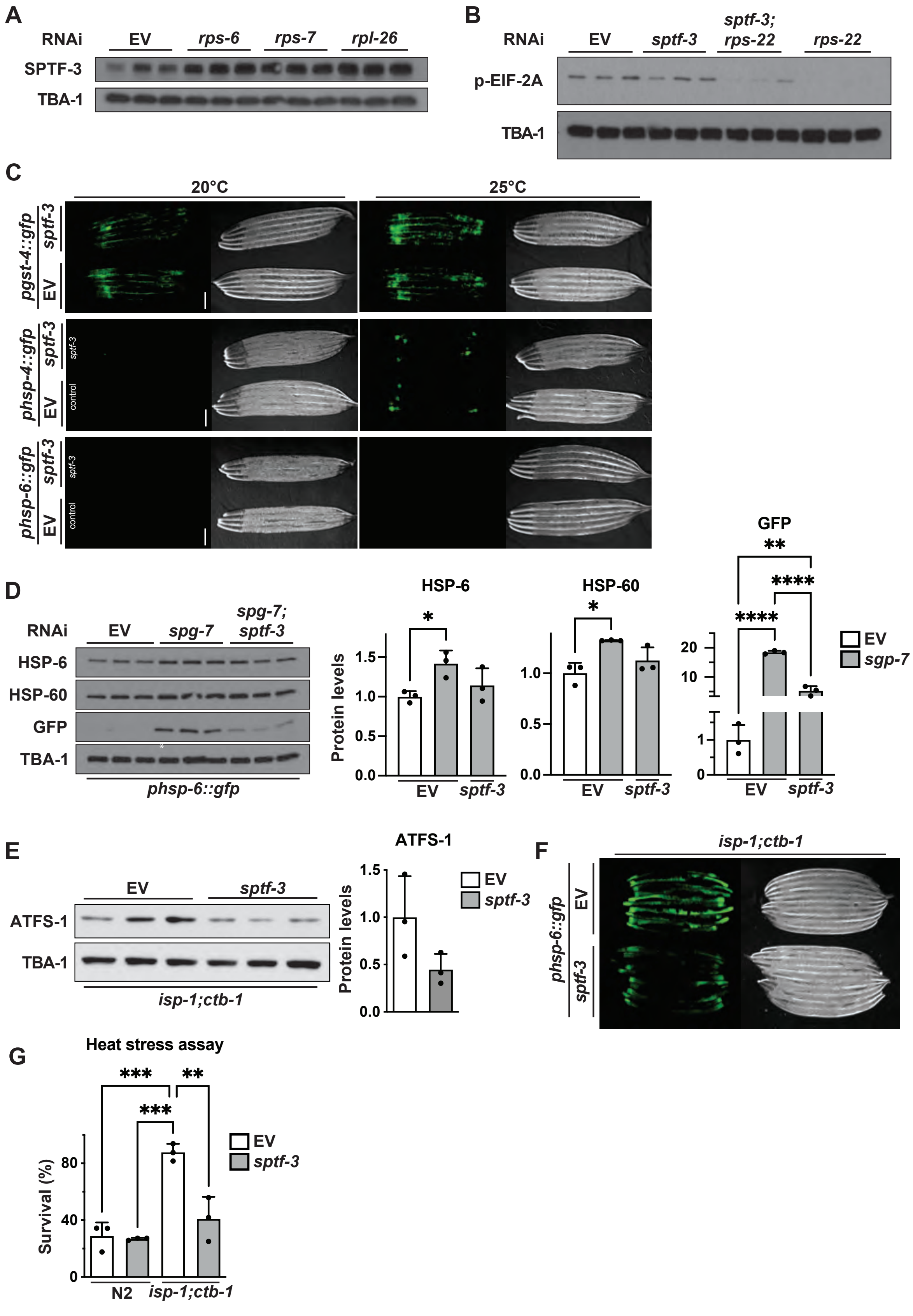

**Extended Data Fig.4 SPTF-3 is required for stress adaptation.**

**A)** Western blot analysis of SPTF-3 levels upon knockdown of ribosomal subunits *rps-6, rps-7* or *rpl-26*; **B)** Steady state levels of phosphorylated (p-)eIF-2α upon *sptf-3, rps-22* or combined *sptf-3/rps-22* RNA; **C)** Fluorescent images of stress markers upon *sptf-3* RNAi on either 20°C or 25°C. Oxidative stress was shown by *pgst-4::gfp* reporter strain, endoplasmic reticulum stress by *phsp-4::gfp* reporter strain and $UPR^{mt}$ - mitochondrial unfolded protein response by *phsp-6::gfp* reporter strain; **D)** Steady-state levels of $UPR^{mt}$ proteins HSP-6, HSP-60 and HSP-6::GFP upon *spg-7,* combined *spg-7/sptf-3* RNAi in animals expressing *phsp-6::gfp.* Western blot (*left*) and analysis (*right*) are shown; **E)** Western blot analysis of ATSF-1 upon *sptf-3* RNAi in *isp-1(qm150)* IV; *ctb-1(qm189)* animals. Western blot (*left*) and quantitative analysis (*right*); **F)** *hsp-6* expression levels measured using *phsp-6::gfp* reporter upon *sptf-3* RNAi in *isp-1(qm150)* IV; *ctb-1(qm189)* mutant; **G)** Stress resistance analysed by heat stress assay upon *sptf-3* RNAi in *isp-1(qm150)* IV; *ctb-1(qm189)* mutant. Data are presented as mean ± SD. $^{*}p<0.05$, $^{**}p<0.01$, $^{***}p<0.001$, $^{****}p<0.0001$. **D, G)** one-way ANOVA with Tukey post hoc test. **E)** unpaired t-test.

#### Extended Data Fig.5

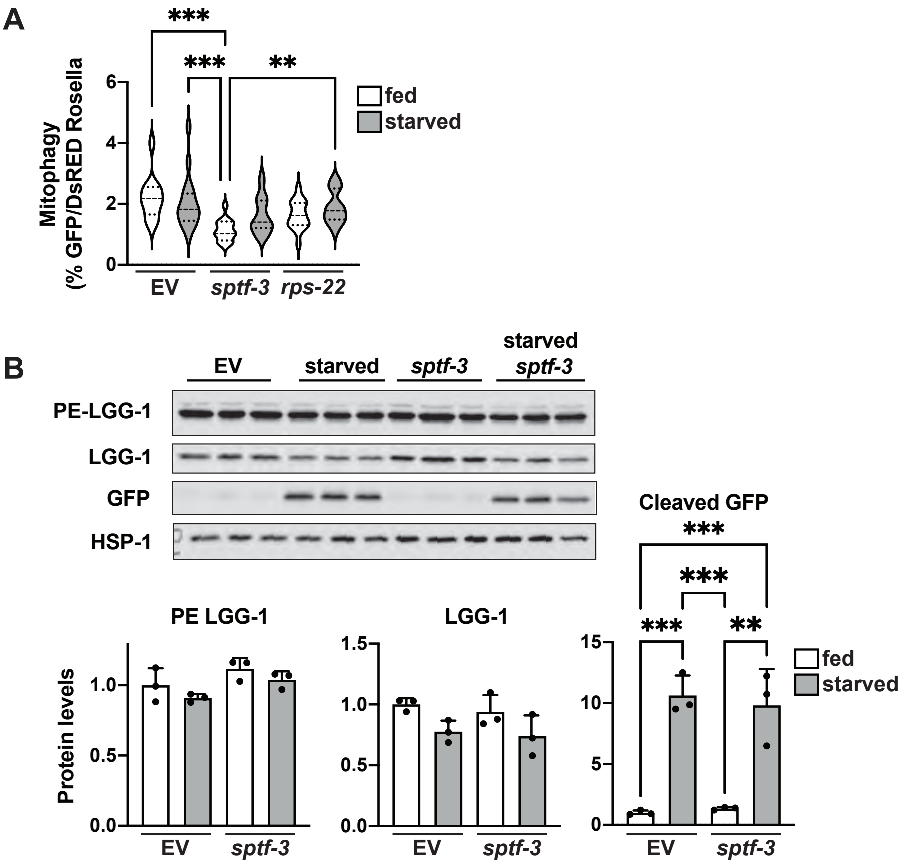


**Extended Data Fig.5 SPTF-3 pathway is autophagy-independent.**

**A)** Mitophagy was measured using the Rosella biosensor[57, 58] by analysing the ratio pH-sensitive GFP versus pH-insensitive DsRed in animals exposed to *sptf-3* and *rps-22* RNAi upon 3h starvation; **B)** Steady-state autophagy protein levels PE-LGG-1, LGG-1 and cleaved GFP levels were measured in control and *sptf-3(n4850)* mutant upon 3h starvation. Western blot (*up*) and analysis (*down*) are shown. Loading control was HSP-1. Data are presented as mean ± SD. $^{**}p<0.01$, $^{***}p<0.001$, one-way ANOVA with Tukey post hoc test.

**Extended Data Table 1 - 4**

| Gene | GFP signal | Gene | GFP signal | Gene | GFP signal |
|---|---|---|---|---|---|
| control | → | *hum-5* | ↗ | *hpl-2* | ↗ |
| *rps-0* | ↑↑ | *atp-2* | ↑ | *Y47D3A.29* | ↗ |
| *rps-12* | ↑↑ | *cdtl-7* | ↗ | *rabx-5* | ↗ |
| *rps-1* | ↑↑ | *rpc-2* | ↑ | *ubq-2* | ↗ |
| *rps-22* | ↑↑ | *polq-1* | ↗ | *rpt-6* | ↗ |
| *rpl-9* | ↑↑ | *him-10* | ↗ | *ccf-1* | ↗ |
| *rpl-35* | ↑↑ | *F01F1.11* | ↗ | *mrg-1* | ↗ |
| *rpl-3* | ↑↑ | *mig-21* | ↗ | *T27E9.2* | ↗ |
| *C34C12.8* | ↗ | *pars-1* | ↗ | *ubl-1* | ↗ |
| *T24C4.5* | ↗ | *ubq-1* | ↗ | *ral-1* | ↗ |
| *W06E11.1* | ↗ | *hmg-4* | ↗ | | |

**Extended Data Table 1 Candidates from suppressor screen using animals expressing *pmtss-1::gfp***

| Gene | GFP | Gene | GFP | Gene | GFP |
|---|---|---|---|---|---|
| Control | → | *rps-2* | ↑↑ | eIF3h | → |
| Ribosome | | *rps-20* | ↑↑ | eIF3j | → |
| *rpl-1* | ↑↑ | *rps-21* | ↑↑ | eIF3k | → |
| *rpl-10* | ↑↑ | *rps-22* | ↑↑ | eIF4A | → |
| *rpl-11.1* | ↑↑ | *rps-23* | ↑↑ | eIF4A-p56 | → |
| *rpl-11.2* | ↑↑ | *rps-24* | ↑↑ | eIF4A-47 | → |
| *rpl-12* | ↑↑ | *rps-26* | ↑↑ | eIF4A-DDX19 | → |
| *rpl-13* | ↑↑ | *rps-27* | ↑↑ | eIF4A-Prp5 | → |
| *rpl-14* | ↑↑ | *rps-28* | ↑↑ | eIF4E-1 | → |
| *rpl-16* | ↑↑ | *rps-29* | ↑↑ | eIF4E-2 | → |
| *rpl-17* | ↑↑ | *rps-30* | ↑↑ | eIF4E-3 | → |
| *rpl-18* | ↑↑ | *rps-4* | ↑↑ | eIF4E-4 | → |
| *rpl-19* | ↑↑ | *rps-5* | ↑↑ | eIF4G | → |
| *rpl-2* | ↑↑ | *rps-7* | ↑↑ | eIF5 | → |
| *rpl-20* | ↑↑ | *rps-8* | ↑↑ | eIF5A | → |
| *rpl-21* | ↑↑ | *rps-9* | ↑↑ | eIF5A | → |
| *rpl-23* | ↑↑ | *rpl-15* | → | eIF5B | → |
| *rpl-24.1* | ↑↑ | *rpl-24.2* | → | Translation elongation factor | |
| *rpl-26* | ↑↑ | *rpl-25.2* | → | eEF1A | → |
| *rpl-27* | ↑↑ | *rpl-28* | → | eEF1A | → |
| *rpl-3* | ↑↑ | *rpl-29* | → | eEF1B | → |
| *rpl-30* | ↑↑ | *rpl-4* | → | eEF1B | → |
| *rpl-32* | ↑↑ | *rpl-42* | → | eEF2 | → |
| *rpl-33* | ↑↑ | *rps-15* | → | eEF2 | → |
| *rpl-34* | ↑↑ | *rps-25* | → | Translation termination factor | |
| *rpl-35* | ↑↑ | *rps-3* | → | eRF1 | → |
| *rpl-36* | ↑↑ | Translation initiation factor | | eRF3 | → |
| *rpl-37* | ↑↑ | eIF1 | → | Translational control | |
| *rpl-38* | ↑↑ | eIF1A | → | TAP42 | → |
| *rpl-41* | ↑↑ | eIF2A | → | SIT4.1 | → |
| *rpl-5* | ↑↑ | eIF2α | → | SIT4.2 | → |
| *rpl-6* | ↑↑ | eIF2α β | → | Akt | → |
| *rpl-7* | ↑↑ | eIF2B | → | PERK | → |
| *rpl-7A* | ↑↑ | eIF2Bα | → | MKK7/ JNKK2 | → |
| *rpl-9* | ↑↑ | eIF2Bβ | → | MKK4 | → |
| *rps-0* | ↑↑ | eIF2Bε | → | MAPK7 | → |
| *rps-1* | ↑↑ | eIF2C-1 | → | MAPK7 | → |
| *rps-10* | ↑↑ | eIF2C-2 | → | ERN1/ IRE1 | → |
| *rps-11* | ↑↑ | eIF3a | → | TOR | → |
| *rps-12* | ↑↑ | eIF3a | → | PI3K like | → |
| *rps-13* | ↑↑ | eIF3b | → | PI4K | → |
| *rps-14* | ↑↑ | eIF3c | → | PI4K | → |
| *rps-16* | ↑↑ | eIF3d | → | Raptor | → |
| *rps-17* | ↑↑ | eIF3e | → | GCN-1 | → |
| *rps-18* | ↑↑ | eIF3f | → | GCN-2 | → |
| *rps-19* | ↑↑ | eIF3g | → | | |

**Extended Data Table 2 Candidates from suppressor screen focusing on genes related to translation using animals expressing *pmtss-1::gfp.***

| Gene | M82 | N81 | Gene | M82 | N81 | Gene | M82 | N81 |
|---|---|---|---|---|---|---|---|---|
| Respiratory chain subunits | | | *T05H4.12* | + | + | *rpom-1* | + | - |
| *F31D4.9* | + | + | *asg-2* | + | + | *sco-1* | + | + |
| *Y53G8AL.2* | + | + | Mitochondrial ribosomes | | | *mtch-1* | + | + |
| *C34B2.8* | + | + | *mrps-7* | + | + | *atad-3* | + | + |
| *Y18D10A.3* | + | + | *mrps-12* | - | + | *cchl-1* | + | + |
| *C50B8.3* | - | + | *mrps-17* | + | + | *mecr-1* | + | + |
| *Y116A8C.30* | + | + | *mrps-23* | + | + | *tufm-1* | + | + |
| *Y51H1A.3* | + | + | *mrps-34* | + | + | *idhg-1* | + | + |
| *nuo-5* | + | + | *mrpl-9* | - | + | *akap-1* | + | + |
| *nuo-2* | + | + | *mrpl-15* | + | + | *mdh-2* | + | + |
| *nduf-5* | + | - | *mrpl-17* | + | + | *Y66A7A.2* | + | - |
| *W10D5.2* | + | + | *mrpl-20* | + | + | *wah-1* | + | - |
| *nuo-1* | + | + | *mrpl-28* | + | + | *mai-2* | + | + |
| *C03G5.1* | + | + | *mrpl-32* | + | + | *srs-1* | + | - |
| *cyc-1* | + | + | *mrpl-47* | - | + | *ogdh-1* | + | + |
| *cyc-2.1* | + | + | *mrpl-49* | + | + | *acl-3* | + | + |
| *ucr-2.2* | + | + | *mrpl-51* | + | + | *ech-2* | + | + |
| *ucr-2.1* | + | + | *mrpl-53* | - | + | *tgt-1* | + | + |
| *ucr-1* | + | + | *mrpl-55* | + | + | *nkcc-1* | + | - |
| *W09C5.8* | + | + | Other mitochondrial proteins | | | *coq-3* | + | + |
| *cco-2* | + | + | *clpp-1* | + | + | *dhs-13* | + | + |
| *cco-1* | + | + | *ZC376.7* | + | + | *tomm-20* | + | + |
| *tag-174* | - | + | *mtss-1* | + | + | *letm-1* | + | + |
| *H28O16.1* | + | + | *cts-1* | + | + | *pus-1* | + | + |
| *atp-2* | + | + | *Y67H2A.4* | + | + | *immt-1* | + | + |
| *asb-2* | + | + | *tag-61* | + | + | *mai-1* | + | + |
| *Y82E9BR.3* | + | + | *pdp-1* | + | + | *got-2.2* | + | + |
| *Y82E9BR.3* | + | + | *T25G3.4* | + | + | *dct-1* | + | + |
| *Y82E9BR.3* | + | + | *nmat-2* | + | - | *dhb-1* | + | + |
| *atp-3* | + | + | *H25P06.1* | + | - | | | |
| *R04F11.2* | + | + | *timm-23* | | | | | |

**Extended Data Table 3 Transcriptional targets of SPTF-3.**

(+) or (-) indicates precipitation status of the genomic region with either M82 or N81 SPTF-3 antibody [21]

| Genotype | Condition | Mean±SE | Median survival | p value (Log-rank test) | Number of animals died/total |
|---|---|---|---|---|---|
| Fig 4F | 25°C | | | | |
| N2 | EV[a] | 13.79±0.349 | 14 | | 89/100 |
| N2 | *sptf-3*[b] | 7.24±0.173 | 7 | < 0.0001[a] | 58/61 |
| *isp-1;ctb-1* | EV[b] | 19.20±0.668 | 22 | < 0.0001[a]<br>< 0.0001[b] | 80/100 |
| *isp-1;ctb-1* | *sptf-3* | 7.09±0.260 | 6 | < 0.0001[a]<br>ns[b]<br>< 0.0001[c] | 75/100 |
| Fig 4G | 20°C | | | | |
| N2 | EV[a] | 22.39±0.295 | 21 | | 173/200 |
| N2 | *rps-22*[b] | 19.95±0.358 | 20 | ns[a] | 174/200 |
| *sptf-3 (n4850)* | EV[c] | 20.63±0.306 | 20 | ns[a]<br>ns[b] | 181/200 |
| *sptf-3 (n4850)* | *rps-22* | 16.71±0.343 | 15 | < 0.0001[a]<br>< 0.0001[b]<br>< 0.0001[c] | 184/200 |
| Fig 5G | 20°C | | | | |
| N2 | fed[a] | 22.39±0.295 | 21 | | 173/200 |
| N2 | starved[b] | 17.56±0.256 | 16 | < 0.0001[a] | 189/200 |
| *sptf-3 (n4850)* | fed[c] | 20.63±0.306 | 20 | ns[a]<br>< 0.0001[b] | 181/200 |
| *sptf-3 (n4850)* | starved | 14.57±0.240 | 15 | < 0.0001[a]<br>< 0.0001[b]<br>< 0.0001[c] | 187/200 |

**Extended Data Table 4 Lifespan statistics.**

[a, b, c] in column p-value is referring to other conditions of the same figure as stated in column condition.
